# Dynamic control of pathway expression with riboregulated switchable feedback promoters

**DOI:** 10.1101/529180

**Authors:** Cameron J. Glasscock, John T. Lazar, Bradley W. Biggs, Jack H. Arnold, Min Kyoung Kang, Danielle Tullman-Ercek, Keith Tyo, Julius B. Lucks

## Abstract

Dynamic pathway regulation has emerged as a promising strategy in metabolic engineering for improved system productivity and yield, and continues to grow in sophistication. Bacterial stress-response promoters allow dynamic gene regulation using the host’s natural transcriptional networks, but lack the flexibility to control the expression timing and overall magnitude of pathway genes. Here, we report a strategy that uses RNA transcriptional regulators to introduce another layer of control over the output of natural stress-response promoters. This new class of gene expression cassette, called a riboregulated switchable feedback promoter (rSFP), can be modularly activated using a variety of mechanisms, from manual induction to quorum sensing. We develop and apply rSFPs to regulate a toxic cytochrome P450 enzyme in the context of a Taxol precursor biosynthesis pathway and show this leads to 2.4x fold higher titers than from the best reported strain. We envision that rSFPs will become a valuable tool for flexible and dynamic control of gene expression in metabolic engineering, protein and biologic production, and many other applications.

## Introduction

Sustainable production of chemicals and materials in microbes through metabolic engineering^1,2^ is a long-standing focus of synthetic biology. A primary challenge in metabolic engineering is the burden and toxicity on engineered cells owing to heterologous enzyme expression and unnecessary intracellular accumulation of toxic pathway intermediates^3,4^. This deleterious effect to the host often results in a loss to productivity and yield, creating a pressing need for strategies that can alleviate or avoid these pitfalls. This is nontrivial because each pathway can present unique cellular stresses, making it difficult to find generalizable solutions.

Synthetic biologists have sought to alleviate pathway toxicity by using dynamic pathway regulation to precisely tune the level and timing of enzyme expression^5–8^. These systems are designed to adaptively adjust enzyme expression in response to changes in growth phase, cellular stress, fermentation conditions, and pathway intermediate concentrations so that they maintain an optimal concentration of enzymes that can vary over time. In order to implement these designs, synthetic biologists have created synthetic feedback networks that dynamically control gene expression using regulatory parts, such as engineered transcription factors^9–11^ or ligand-induced ribozymes^12^, that respond to relevant cues.

While these systems represent important advances, synthetic feedback networks are often difficult to construct because the sensors required for specific inputs are hard to design or source from nature, and the added burden of expressing regulatory components can itself negatively impact the host^13^. Ultimately, this means that synthetic feedback networks require considerable additional engineering to match the specific requirements of every application. On the other hand, nature has evolved stress-responsive feedback networks that are already compatible with host cells. This is a result of the fact that microbial cells persist and thrive in changing environments due to their ability to sense and respond to stresses and environmental conditions^14^. Much of this ability is encoded within regulatory elements called stress-response promoters that integrate signals from complex and interconnected transcriptional networks to modulate mRNA synthesis in response to specific cellular stresses^15^. This creates the possibility of using stress-response promoters to regulate heterologous pathway expression as a means to implement genetic feedback networks that lead to improvements in productivity and yield. In fact, synthetic biologists have used stress-response promoters to control pathway expression, leading to notable improvements to productivity and yield for protein expression^16^ and industrially important pathways, such as the artemisinin precursor amorphadiene^17^ and n-butanol^18^. However, stress-response promoters have not been widely adopted, as their complexity makes it difficult to fine-tune their behavior for specific applications. As their function is determined by the complex topologies of natural genetic networks, there are no simple methods to tune either the timing or overall magnitude of their transcriptional outputs – two key parameters that are important for optimizing metabolic pathway productivity and yield^19^.

To address this limitation, we sought to create a new regulatory motif called a switchable feedback promoter (SFP) that combines the feedback properties of natural stress-response promoters with regulators that offer control of the timing and overall magnitude of transcriptional outputs (Figure 1A-C). The SFP concept is general, and can be implemented in several ways including engineering transcription factor operator sites within the stress-response promoter region^20^. However, the architecture of many stress-response promoters is still unknown, making the rational design of transcription factor-based SFPs difficult. Instead, we utilized *trans*-acting synthetic RNA regulators^21,22^, which can be configured to control transcription in the case of small transcription activating RNAs (STARs)^21^, or translation in the case of toehold switches^22^. The key feature of both STARs and toehold switches is that they have well-defined composition rules such that they can be inserted into a gene expression construct without modification or disruption of the desired promoter sequence. In this way, riboregulated SFPs (rSFPs) can be created by inserting a STAR or toehold target binding region 3’ of a natural stress-response promoter (Fig. 1B). By default, when transcribed, these RNA sequences fold into structures that block gene expression. However, binding of the trans-acting STAR or toehold trigger RNA changes these structures to allow gene expression. This enables the timing and overall magnitude of the rSFP output to be controlled with any strategy that can regulate the expression of the *trans*-acting RNA (Fig. 1C). These regulators have additional advantages that make inclusion in rSFPs promising, including the availability of large libraries of orthogonal STARs and toehold switches with a range of functional properties (ON/OFF levels)^22,23^ and their having compact DNA footprints (<100nts). Furthermore, they have been shown to control gene expression in a variety of contexts, including within metabolic pathways^23^.

**Figure 1.**
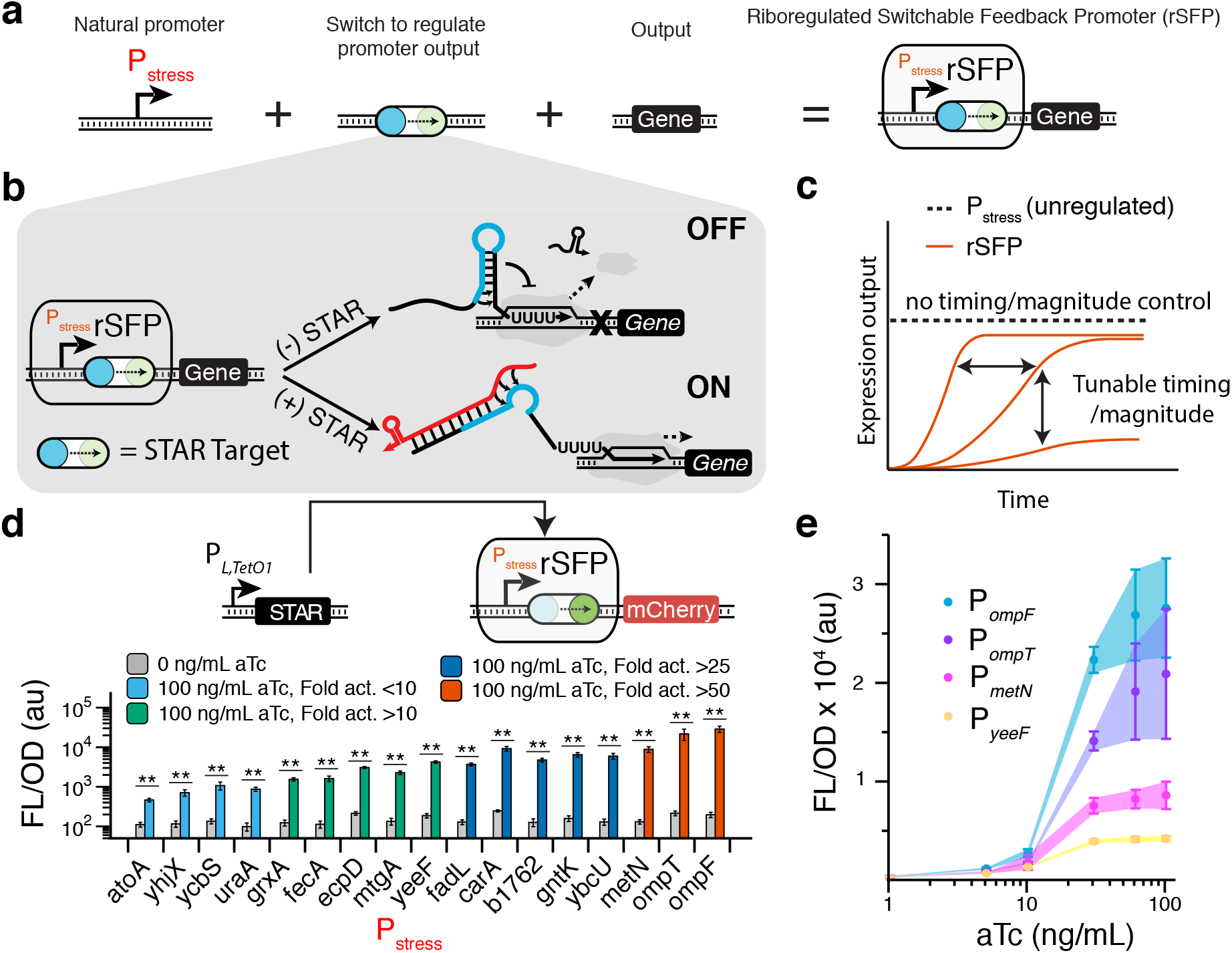
Riboregulated switchable feedback promoters. (**a**) Riboregulated switchable feedback promoters (rSFPs) are composed of a natural stress-response promoter and an RNA transcriptional switch that allows control over the output of native stress-mediated transcriptional networks. (**b**) Schematic of the small transcription activating RNA (STAR) transcriptional switch mechanism used in rSFPs. A Target DNA sequence (switch symbol) is placed 3’ of a stress-response promoter. The transcribed Target RNA is designed to fold into an intrinsic transcription terminator hairpin, composed of a hairpin structure followed immediately by a poly uracil sequence. The formation of this terminator hairpin causes RNA polymerase (RNAP) to terminate transcription upstream of the gene to be regulated (gene OFF). A separately transcribed STAR RNA (colored red) can bind to both the linear region and the 5’ half of the terminator hairpin (colored blue) of the Target RNA, preventing its formation and allowing transcription elongation of the downstream gene (gene ON). In this way the output of the stress-response promoter is controlled by STAR expression, which adds an additional layer of regulation to that present within the stress-mediated transcriptional network that governs expression at the stress-response promoter. (**c**) Illustration of expression control enabled by rSFPs. Natural stress-response promoters (dashed line) can exhibit dynamic behaviors in response to stress but are fixed with regards to user-defined timing and overall expression magnitude. rSFPs (red lines) use the additional layer of regulation to resolve this issue and allow control of timing and overall expression magnitude by gating transcriptional output with a *trans*-acting RNA regulatory switch. (**d**) Characterization of rSFP variants containing unique envelope stress-response promoters. *P_L,TetO1_* inducible STAR expression is used to activate rSFPs containing a natural stress-response promoter upstream of a STAR Target sequence, a ribosome binding site, and a red fluorescent protein (mCherry) coding sequence. Fluorescence characterization was performed on *E. coli* transformed with plasmids encoding each rSFP controlling mCherry expression in the absence and presence of 100 ng/mL aTc. (**e**) rSFPs enable titration of natural stress-response promoter output. Fluorescence characterization performed on *E. coli* cells containing rSFPs controlling mCherry expression under different levels of aTc induction. Data in **d, e** represent mean values in units of arbitrary fluorescence/optical density (FL/OD) and error bars represent s.d. of at least n = 7 biological replicates. * indicate a statistically significant difference in FL/OD by a two-tailed Welch’s t-test (* = P < 0.05, ** = P < 0.005).

Here we report the creation and characterization of a library of STAR-mediated rSFPs, and their application to optimizing the yield of a metabolic pathway that produces an oxygenated taxane precursor to the anticancer drug Taxol. We first show that we can create a library of 17 rSFPs by interfacing STARs with natural *Escherichia coli* stress-response promoters and placing *trans*-acting STAR production under control of an inducible promoter. We then applied rSFPs to control the expression of a plant cytochrome P450 that is known to cause envelope stress^24^ in the context of a pathway that produces an oxygenated Taxol precursor^25^. By screening rSFPs for oxygenated taxane production, we were able to find multiple rSFPs that showed improvement in both overall and oxygenated taxane titers compared to the previously reported best strain. We next used the external control of rSFPs to systematically optimize both timing and expression level to ultimately find pathway conditions that produce 25.4 mg/L of oxygenated taxanes and 39.0 mg/L of total taxanes, representing a 2.4x and a 3.6x fold improvement over the current state-of-the-art, respectively. To demonstrate the use of other control points for rSFPs, we next sought to interface them with a quorum sensing system and show that quorum sensing rSFPs offer completely autonomous pathway expression regulation with yields similar to our fully optimized system without costly external inducers.

Overall, rSFPs are a novel and general strategy to achieve dynamic regulation of metabolic pathway enzymes and we envision them to be broadly useful for introducing controllable stress-response promoters in many synthetic biology applications.

## Results

### Riboregulated switchable feedback promoters (rSFPs) enable tunable outputs from stress-response promoters

We chose to build rSFPs with STARs because they exhibit low leak and high dynamic range comparable to exemplary protein-based regulators^23^. STARs activate transcription by disrupting the folding pathway of a terminator hairpin sequence, called a Target, that is placed upstream of the gene to be regulated (Fig. 1B). In the absence of a STAR, the Target region folds into an intrinsic terminator hairpin which stops transcription before reaching the downstream gene. When present, a STAR RNA can bind to the 5’ portion of the terminator hairpin, preventing its formation, and allowing transcription. rSFPs are then created by inserting a Target sequence 3’ of a candidate stress-response promoter. In this way, the introduction of the STAR/Target adds an additional layer of control to the stress-response promoter, effectively gating its transcriptional output through the additional regulation of STAR RNA expression, which can be controlled using a variety of mechanisms, including manual inducible promoters or quorum-sensing systems.

Our initial rSFP designs utilized a previously developed STAR^23^ under the well-characterized inducible system TetR/P_*L,TetO1*_^20^ interfaced with a library of 17 putative membrane stress-responsive promoters^17,18^. These promoters were chosen as several had been previously identified to regulate a biofuel transporter protein in *E. coli*^18^, and could be valuable for dynamic regulation of membrane proteins in metabolic pathways. To construct and characterize these rSFPs, a STAR Target sequence was cloned immediately 3’ of each promoter to regulate expression of an mCherry reporter, and its cognate STAR was cloned in a second P_*L,TetO1*_ plasmid. Plasmids were transformed into *E. coli* and fluorescence was measured with and without the presence of the P_*L,TetO1*_ inducer anhydrotetracycline (aTc) at saturating levels (100 ng/mL). We found that induction of P_*L,TetO1*_-STAR resulted in significant activation from all members of the stress-response promoter library (Fig. 1D), exemplifying the modularity of the rSFP concept. We also observed that 8 library members were activated by greater than 25x fold in the presence of aTc, with a maximum activation of nearly 150x fold (SI Fig. 1). We next selected a set of high-performing rSFPs and characterized their transfer functions by titrating levels of aTc and found that all exhibited a similar transfer function shape, though with different maximal activation levels (Fig. 1E). This is evidence that the transfer function of the P_*L,TetO1*_ regulatory system can be overlaid on a range of stress response promoters through the STAR intermediate. Overall these results demonstrated that we can create a library of rSFPs that provide tunable control of gene expression level by selecting different stress-response promoters and manipulating inducer concentration.

### rSFPs enhance production of an oxygenated Taxol precursor

We next tested the ability of rSFPs to regulate expression of a challenging metabolic pathway enzyme. As a model system, we chose a portion of the anticancer drug Paclitaxel’s biosynthesis pathway that has been previously reconstituted in *E. coli*^24^. Specifically, we focused on the first P450-mediated step where taxadiene is oxygenated by the membrane anchored cytochrome P450 CYP725A4 (Fig. 2A). This system is an ideal test bed for the use of rSFPs because CYP725A4 expression causes membrane stress due to lipid anchoring of an N-terminal domain. This stress appears to reduce pathway productivity and makes pathway optimizations extremely difficult^24^. The sensitivity of product titers to expression level of CYP725A4 must be carefully managed, as too low expression will create a bottleneck in oxygenated taxane synthesis, but too high expression will also suppress synthesis due to stress. Previous reports to optimize this system required significant experimental effort^24^, exemplifying the importance of new pathway engineering strategies, but also providing a competitive benchmark for comparison with rSFPs. Furthermore, this problematic expression is not unique to CYP725A4, but extends to many P450’s^26,27^, along with other classes of proteins such as transporters^18^ and glycosylation enzymes^28,29^. Therefore, the system is a model challenge for testing the concept of using rSFPs to leverage external control of natural stress-response promoters to maintain pathway expression in a narrow optimal range.

**Figure 2.**
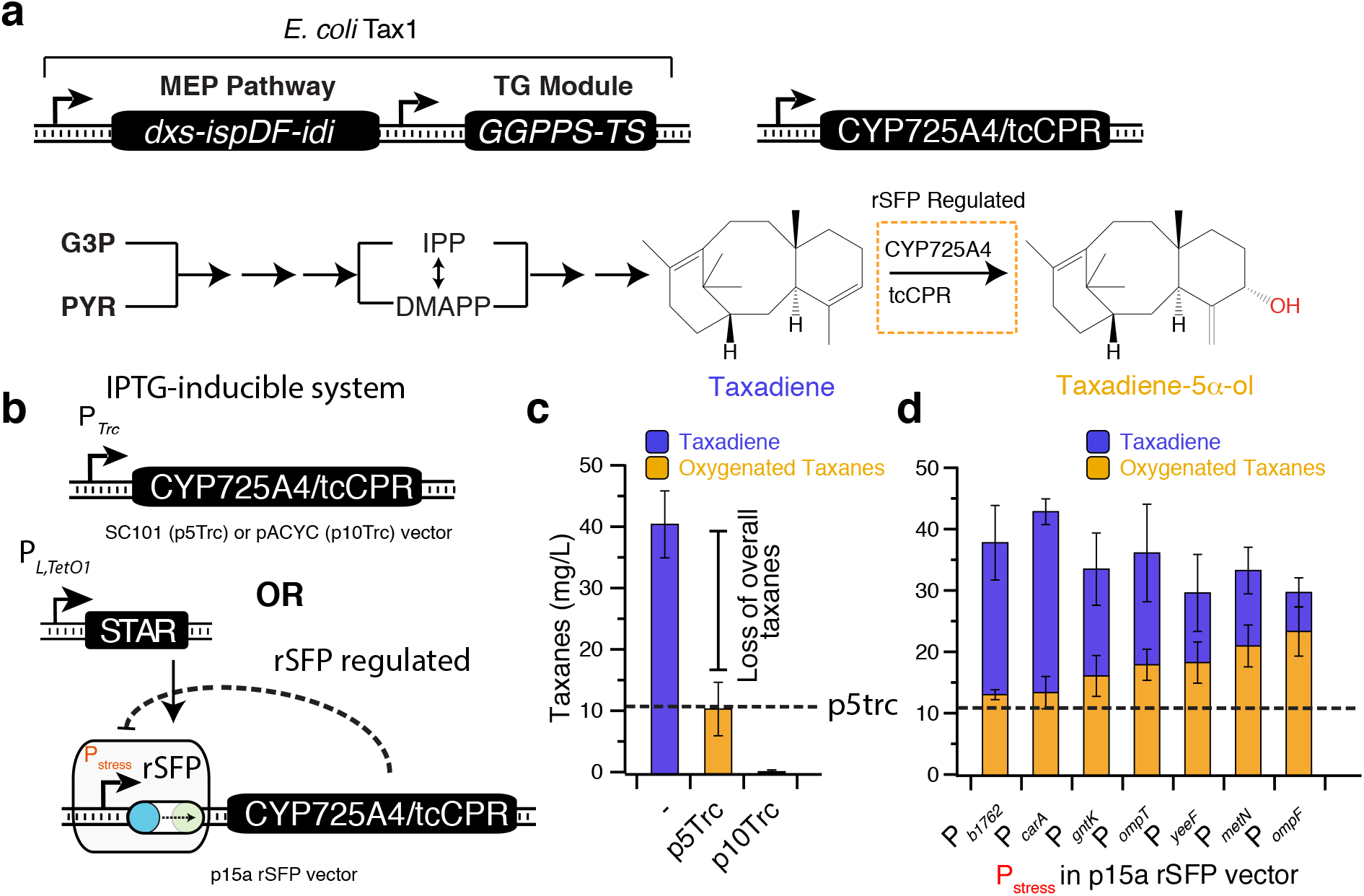
rSFPs enhance productivity of a Taxol precursor synthesis pathway in *E. coli*. (**a**) Taxol biosynthesis schematic depicting an abbreviated overview of the Taxol precursor pathway involving the toxic cytochrome P450 725A4 (CYP725A4) enzyme. In the *E. coli* strain Tax1, the methylerythritol phosphate (MEP) pathway and taxadiene synthase/geryanlgeranyl diphosphate (GGPP) synthase (TG) module convert glyceraldehyde-3-phosphate (G3P) and pyruvate (PYR) into the 20-carbon backbone taxa-4 (5),11 (12)-diene (taxadiene). Taxadiene is oxygenated by the membrane-anchored CYP725A4 fused with its reductase partner (tcCPR) to form taxadiene-5α-ol. rSFPs utilizing envelope stress-response promoters are applied to control the expression of CYP725A4/tcCPR. IPP = isopentenyl diphosphate, DMAP = dimethylallyl diphosphate. (**b**) Plasmids used for CYP725A4/tcCPR expression in *E. coli* Tax1. CYP725A4/tcCPR is expressed from a standard IPTG-inducible P*_Trc_* promoter in either a low copy (p5Trc, SC101) or medium copy (p10Trc, pACYC) plasmid, or from rSFPs with various P_stress_ promoters encoded on a p15a plasmid. P_*L,TetO1*_-STAR was used to activate rSFP expression. rSFPs allow CYP725A4/tcCPR expression to be controlled by externally supplied aTc and feedback-regulated by the natural stress response pathways. p5Trc is a gold standard CYP725A4/tcCPR expression system from previously reported optimization efforts. (**c**) Titers of fermentations with empty *E. coli* Tax1 and *E. coli* Tax1 containing low copy p5Trc or medium copy p10Trc expression of CYP725A4/tcCPR. Addition of p5Trc to *E. coli* Tax1 enables production of oxygenated taxanes at a cost to overall taxane production, while addition of p10Trc eliminates nearly all taxane production presumably due to the toxicity of CYP725A4/tcCPR expression. (**d**) Titers of fermentations with *E. coli* Tax1 containing CYP725A4/tcCPR under control of high performing rSFPs. Dashed line represents production of oxygenated taxanes from p5Trc. The P_*ompF*_ rSFP resulted in ~2.2x fold greater oxygenated taxanes (~23.5 mg/L) and ~2.8x fold greater overall taxanes (29.8 mg/L) than the p5Trc strain. Data in **c, d** represent mean values of fermentation titers after 96 hrs and error bars represent s.d. of at least n = 5 biological replicates.

Previous work has shown that expression level of a CYP725A4/tcCPR reductase fusion is critical to achieving high titers of oxygenated taxanes in *E. coli*^24^. A previously optimized low-copy expression vector (p5Trc-CYP725A4/tcCPR) (Fig. 2B) transformed into the *E. coli* Tax1 strain containing genomic modifications to maximize the synthesis of the taxadiene precursor, produces ~11 mg/L of oxygenated taxanes, albeit with a loss to total taxane production. However, increasing expression of the enzyme using a medium copy expression vector (p10Trc) does not increase titer, but causes a complete loss of pathway productivity (Fig. 2C), presumably due to the enzyme’s membrane stress crossing a critical threshold and triggering a global response.

We hypothesized we could achieve greater pathway productivity over the p5Trc benchmark strain by identifying putative envelope stress rSFPs for control of CYP725A4/tcCPR. To test this, the CYP725A4/tcCPR coding sequence was introduced into each one of the 17 rSFP constructs. *E. coli* Tax1 was transformed with each rSFP construct and the P_*L,TetO1*_-STAR plasmid and each tested in the context of taxadiene oxygenation fermentations with addition of 100 ng/mL aTc at inoculation. Using this approach, we found that several performed well against the p5Trc benchmark strain (SI Fig. 2). In particular, 7 of the rSFPs had greater titers of oxygenated taxanes than the p5Trc strain (Fig. 2D), with all also improving overall taxane production. Furthermore, the P_*ompF*_ rSFP resulted in ~2.2x fold greater oxygenated taxanes (~23.5 mg/L) and ~2.8x fold greater overall taxanes (29.8 mg/L) than the p5Trc strain, clearly showing the benefit of rSFP pathway regulation.

To confirm that rSFPs can indeed be feedback regulated by CYP725A4/tcCPR stress, we performed fluorescence analysis of *E. coli* cells containing plasmids for rSFP expression of an mCherry reporter with the top two performing stress-response promoters and the p10Trc plasmid separately expressing CYP725A4/tcCPR, in order to monitor changes in rSFP expression caused by membrane stress (SI Fig. 3A). We observed reduced expression from P_*ompF*_ when p10Trc was present in place of an empty vector (SI Fig. 3B), suggesting that it is indeed responsive to CYP725A4/tcCPR induced stress. On the other hand, a constitutive promoter control had no response as expected. Interestingly, the P_*metN*_ rSFP did not exhibit a reduced expression in response to CYP725A4/tcCPR expression, indicating that not all rSFPs respond to stresses in the same way, as we expected for a diverse set of natural stress-response promoters.

**Figure 3.**
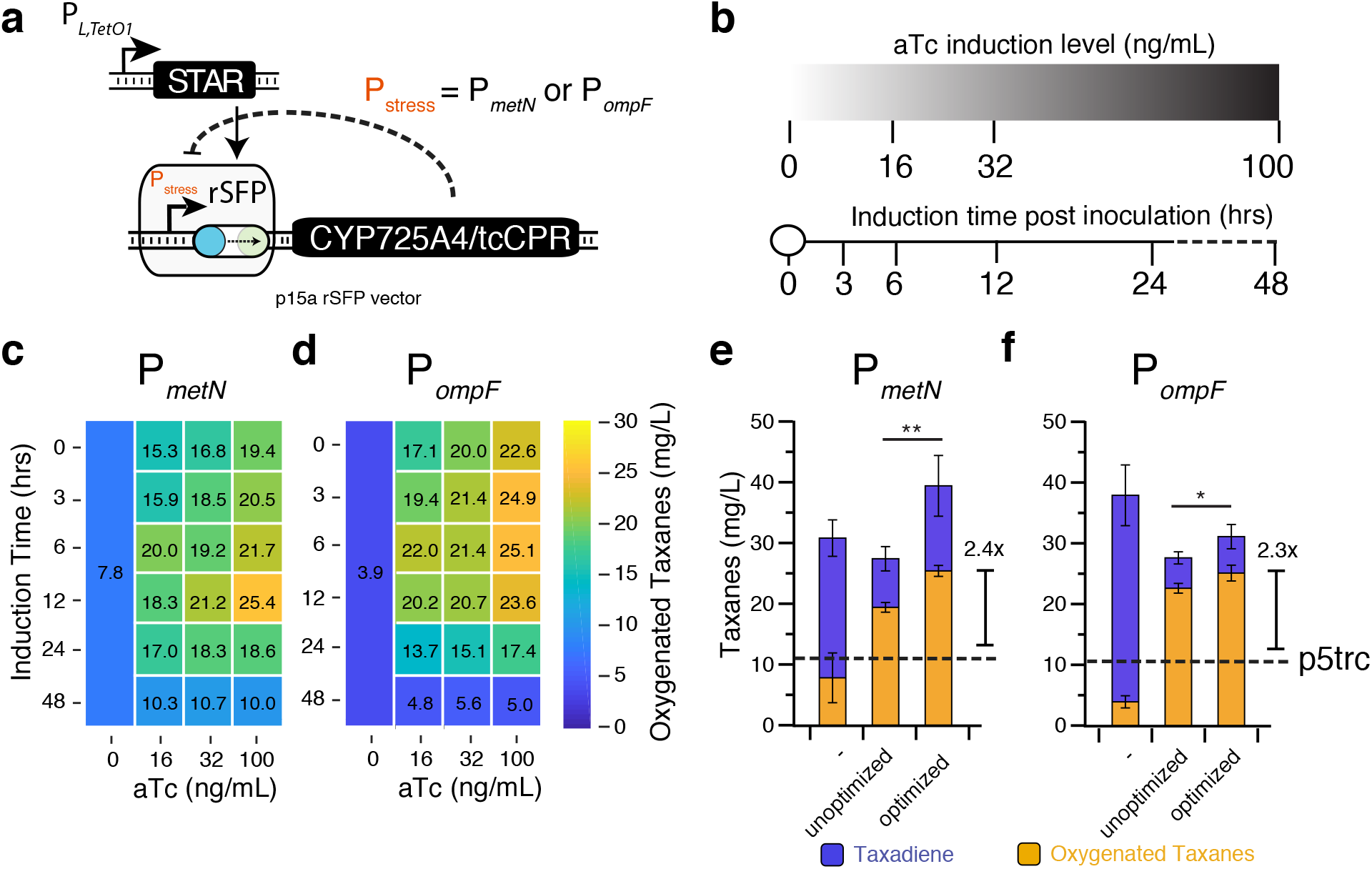
External control of rSFPs enable optimization of induction level and timing from stress-response promoters. (**a**) Plasmids used for CYP725A4/tcCPR expression from rSFPs with P_stress_ promoters encoded on a p15a plasmid in *E. coli* Tax1. P_*L,TetO1*_-STAR was used to activate rSFP expression. (**b**) Conditions of aTc induction level and timing used for rSFP CYP725A4/tcCPR expression optimization. (**c, d**) Induction level and timing optimization of fermentations with *E. coli* Tax1 containing the P_*metN*_ (**c**) or P_*ompF*_ (**d**) rSFP controlling CYP725A4/tcCPR. Heatmap shows the average oxygenated taxane titers for different combinations of aTc concentration and time of induction. (**e, f**) Titers of fermentations with *E. coli* Tax1 containing the P_*metN*_ (**e**) or P_*ompF*_ (**f**) rSFP before (100 ng/mL aTc at 0 hrs) and after induction optimization. Dashed line represents production of oxygenated taxanes from p5Trc. Data in **c, d, e, f** represent mean values of fermentation titers after 96 hrs and error bars represent s.d. of at least n = 3 biological replicates. * indicate a statistically significant difference in oxygenated taxane production by a two-tailed Welch’s t-test (* = P < 0.05, ** = P < 0.005).

Overall these results show that rSFPs can be effectively used to optimize overall pathway expression and that they can exhibit the dynamic feedback behaviors of incorporated stress-response promoters.

### rSFPs allow further pathway optimization through the control of expression timing and overall magnitude

Having shown significant improvements in taxadiene oxygenation with rSFPs, we next sought to test how the external control offered by rSFPs can be used to further optimize induction level and timing of stress-response promoter activity. To test this, we selected the two best rSFP systems and performed a matrix of aTc induction at four levels (0, 16, 32, and 100 ng/mL aTc), which were added at six different induction times (0, 3, 6, 12, 24, 48 hrs) post fermentation inoculation (Fig. 3A-B). We found that oxygenated taxane production with both rSFPs was indeed sensitive to induction level and timing (Fig. 3C,D) and that late induction of P_*metN*_ and P_*ompF*_ rSFPs could improve final titers of oxygenated taxanes even further to 25.4 and 25.1 mg/L, respectively, and overall taxanes to 39.0 and 31.0 mg/L (Fig. 3E,F), representing an overall 2.4x and 2.3x fold improvement over the previous gold standard benchmark in terms of oxygenated taxanes, and 3.6x and 2.9x fold improvements in terms of overall taxanes. These results demonstrate that rSFPs can be implemented to enable rapid tuning of expression timing and overall magnitude of stress-response promoter output to further enhance fermentation titers.

### Quorum-sensing activated rSFPs allow autonomous regulation of pathway expression

Though inducible systems offer flexibility for screening of optimal induction timing, the cost of inducers can be prohibitive at an industrial scale^30,31^, and several efforts have been carried out to design autonomous means of induction. Quorum-sensing (QS) systems that are activated in a cell-density dependent manner offer one such route to this behavior^32^. QS systems have been used with great utility in metabolic engineering to create a separation of cell growth and pathway production phases without the need for a chemical inducer, and provide a natural means for balancing carbon utilization with biomass production^33–35^. We therefore sought to utilize this strategy within our model pathway by leveraging the modularity of rSFPs to be easily configured to utilize different input systems. Specifically, we chose the P_*Lux*_ promoter that is activated by the LuxR transcriptional activator upon sufficient production of the C6-homoserine lactone (HSL) signaling molecule^36^. We cloned a STAR under control of P_*Lux*_ and integrated an operon with the EsaI HSL synthase^37^ and LuxR into the genome of the *E. coli* Tax1 strain to create the Tax1-QS strain (Fig. 4A). When plasmids encoding the expression of P_*Lux*_-STAR and the P_*metN*_ or P_*ompF*_ rSFPs controlling mCherry expression were transformed into *E. coli* Tax1 or Tax1-QS, we found that activation only occurred in the engineered Tax1-QS strain containing EsaI and LuxR (Fig. 4B). These QS-activated rSFPs produced comparable fold activation to manual induction with P_*L,TetO1*_ and exhibited a time dependent activation.

**Figure 4.**
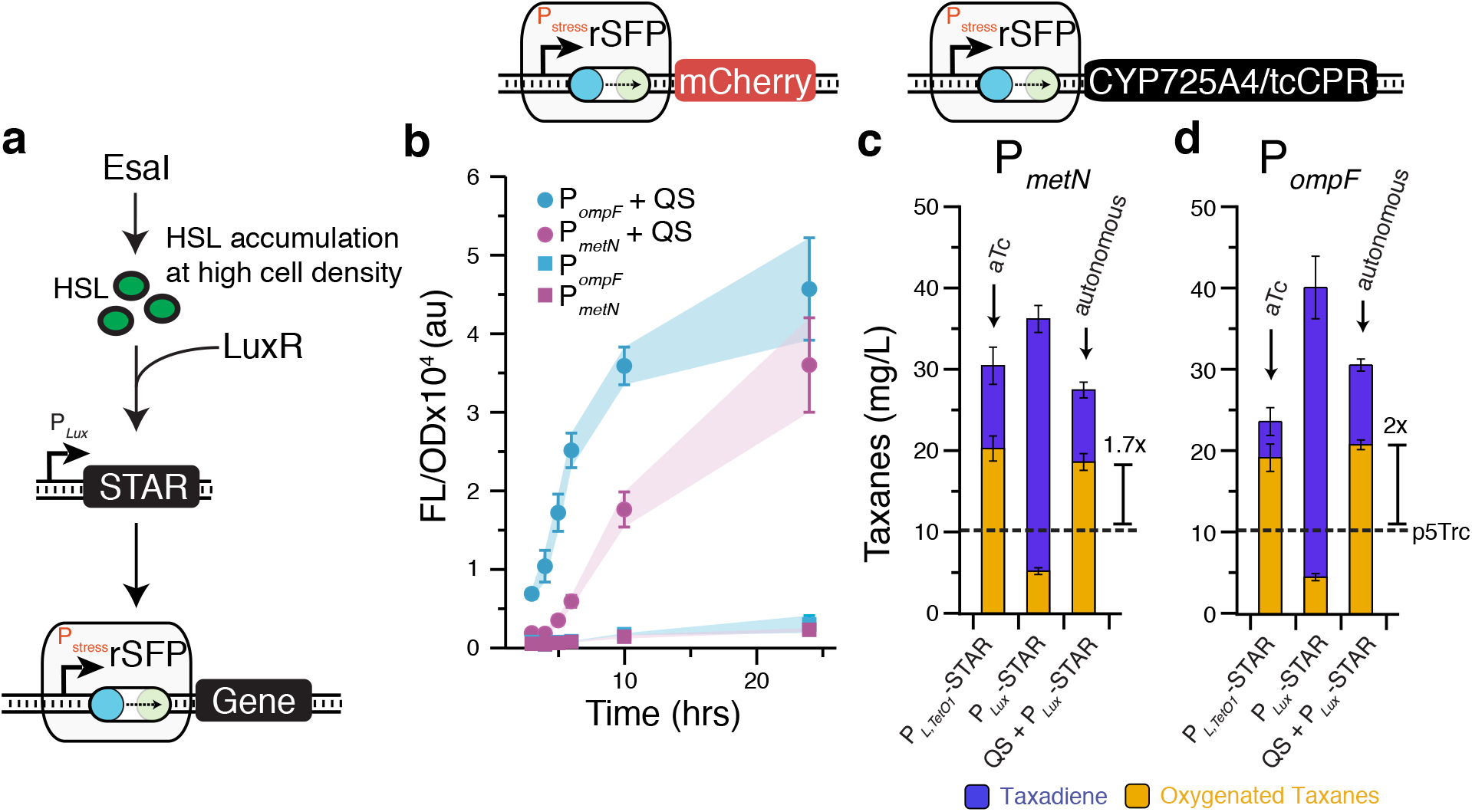
Quorum sensing activation of rSFPs allows autonomous control of CYP725A4 expression. (**a**) Schematic showing quorum-sensing (QS) activation of rSFPs to allow autonomous control of pathway expression. LuxR is activated by C6-HSL produced by the EsaI HSL synthase upon sufficient accumulation due to an increase in cell density. LuxR activation results in STAR production from the P_*Lux*_ promoter, thereby activating rSFP expression. (**b**) Fluorescence experiments showing autonomous activation of P_*ompF*_ and P_*metN*_ rSFPs, configured to control mCherry expression, over time. Fluorescence is only significant with the Tax1-QS strain containing a chromosomal LuxR/EsaI expression cassette, but not with the parent Tax1 strain. (**c, d**) Titers of oxygenated taxadiene fermentations in *E. coli* Tax1 strains containing the P_*metN*_ (**c**) or P_*ompF*_ (**d**) rSFP controlling CYP725A4/tcCPR expression. Left: in *E. coli* Tax1 (without QS insert) containing a P_*L,TetO1*_-STAR plasmid induced by aTc; middle: in *E. coli* Tax1 containing a P_*Lux*_-STAR plasmid; right: in *E. coli* Tax1-QS containing a P_*Lux*_-STAR plasmid activated by EsaI produced HSL. QS-activated rSFPs obtained similar titers to the unoptimized P_*L,TetO1*_-STAR activated rSFPs, but without any external interventions. Dashed line represents production of oxygenated taxanes from p5Trc Fig. 2C. Data in **b** represent mean values in units of arbitrary fluorescence/optical density (FL/OD) and error bars represent s.d. of at least n = 7 biological replicates. Data in **c** represent mean values of fermentation titers after 96 hrs and error bars represent s.d. of at least n = 3 biological replicates.

To demonstrate that QS-activated rSFPs could be used to autonomously control the expression of metabolic pathway enzymes, we applied the P_*metN*_ and P_*ompF*_ QS-activated rSFPs to control the expression of CYP725A4/tcCPR within the taxadiene oxygenation pathway. Fermentations were performed by inoculating cell cultures into media without addition of exogenous inducer. Upon fermentation and analysis, we found that QS-based activation resulted in comparable titers of oxygenated taxanes to those obtained from manual induction of rSFPs with aTc before optimization (Fig. 4C,D). Importantly, this represented 1.7x and 2x fold improvements, respectively, over the previous gold standard, and was achieved with a completely autonomous genetic feedback network without the need for costly inducers.

## Discussion

Here we report the development, characterization and application of switchable feedback promoters that enable an additional synthetic layer of control over natural stress-response promoters. Stress-response promoters are a promising route to achieving dynamic control of heterologous metabolic pathways by acting as sensor-actuators to stresses caused by pathway expression, intermediate metabolites and other fermentation conditions^17,18^. While stress-response promoters can improve production of desired chemicals by regulating expression in response to toxic pathway intermediates and enzymes, they are constrained by their complexity, leading to a lack of control over the timing and overall magnitude of their transcriptional output, which is essential to achieving a separation of growth phase and production phase in large-scale fermentations^38^. By design, the rSFP concept enables this control by introducing an additional regulatory layer within the natural stress-response pathway by gating stress-response promoter outputs with *trans*-acting RNA regulators. The use of an inducible promoter to control RNA regulator synthesis allows modification of the timing and overall magnitude of the natural stress-response promoter outputs. Furthermore, the use of QS systems allows the autonomous activation of rSFPs in a cell-density dependent manner. In this way, rSFPs have modularity both at the lever of their inputs and outputs, and the types of stresses they can respond to through changing of the regulated stress-response promoter. This offers the flexible implementation of controllable stress-response networks in a single compact locus.

In this work, we design and implement rSFPs and demonstrate that they are both modular and tunable – the rSFP concept can be applied to many unique stress-response promoters in a plug-and-play fashion, activator inputs can be easily interchanged, and activated output levels can be modulated by titrating inducer concentrations. Notably, we found that all 17 of the stress-response promoters that were inserted into rSFPs were activated significantly, strongly suggesting that rSFPs can be used with new stress-response promoters as they are discovered, and potentially that the rSFP concept can be used to easily regulate engineered promoter systems as well. These features allow rapid screening of rSFP libraries within combinatorial strain engineering procedures^39^ that could be used by industry to identify effective implementations of dynamic control. In addition, the ability of rSFPs to naturally adapt to an optimal expression level may allow for rapid prototyping of potentially toxic enzymes and pathways without the requisite need to first balance expression levels with constitutive static regulators – speeding the pace of pathway construction for new chemical products.

To demonstrate their utility in the context of optimizing metabolic pathway production, we applied rSFPs to a synthetic Taxol precursor pathway in *E. coli*^25^ by regulating expression of a problematic cytochrome P450 enzyme that causes a membrane stress detrimental to productivity^24^. By screening through a library of envelope-stress-response promoters in rSFPs, we identified variants that improved pathway productivity over a previous strain that had been optimized using a laborious trial-and-error approach. Furthermore, we showed that optimizing rSFP induction timing and magnitude in the fermentation enabled additional improvements, highlighting an advantage of the rSFP system to enable the control of pathway expression timing. We also showed that rSFPs can be controlled by QS systems that do not require addition of an external inducer, enabling fully autonomous control of pathway expression.

Dynamic pathway regulation is a promising strategy in metabolic engineering but can be difficult to implement. The rSFP strategy enables modular and tunable control of endogenous promoters that have evolved sophisticated transcriptional responses to a range of cellular stresses and fermentation conditions. Due to their simplicity, we envision that the rSFP concept will enable streamlined implementation of dynamic regulation into metabolic pathways. Furthermore, given their modularity, we imagine rSFPs will be useful for dynamic control in other applications, such as high-level expression of difficult or toxic proteins, living therapeutics^40^, and cellular diagnostics^41^ where endogenous promoters could be used as sensor-actuators for numerous environments.

## Methods

### Plasmid assembly

All plasmids used in this study can be found in **Supplementary Table 1** with key sequences provided in **Supplementary Tables 2 and 3**. Gibson assembly and inverse PCR (iPCR) was used for construction of all plasmids. All assembled plasmids were verified using DNA sequencing.

### Integration of QS operon into the *E. coli* genome

Strains containing genomic insertions of the EsaI-LuxR operon were created using the clonetegration^42^ platform as summarized in **Supplementary Table 4**. The HK022 plasmid was used to integrate constructs into the *attB* site of the *E. coli* genome. Successful integrations were identified by antibiotic selection and colony PCR according to the published protocol.

### Strains, growth media, *in vivo* bulk fluorescence measurements

Fluorescence characterization experiments for all envelope stress-response promoters were performed in *E. coli* strain Tax1^24^ containing the synthetic pathway for taxadiene biosynthesis or modified Tax1-QS containing the QS operon. Experiments were performed for 7-9 biological replicates collected over three separate days. For each day of fluorescence measurements, plasmid combinations were transformed into chemically competent *E. coli* cells and plated on LB+Agar (Difco) plates containing combinations of 100 μg/mL carbenicillin, 34 μg/mL chloramphenicol and/or 50 μg/mL spectinomycin depending on plasmids used (see SI Table 1 for plasmids used in each experiment), and incubated approximately 17 hours (h) overnight at 37 °C. Plates were taken out of the incubator and left at room temperature for approximately 7 h. Three colonies were used to inoculate three cultures of 300 μL of LB containing antibiotics at the concentrations described above in a 2 mL 96-well block (Costar), and grown for approximately 17 h overnight at 37 °C at 1,000 rpm in a VorTemp 56 (Labnet) bench top shaker. **Figures 1d, 1e**: 4 μL of each overnight culture were added to 196 μL (1:50 dilution) of supplemented M9 minimal media (1 × M9 minimal salts, 1 mM thiamine hydrochloride, 0.4 % glycerol, 0.2 % casamino acids, 2 mM MgSO_4_, 0.1 mM CaCl_2_) containing the selective antibiotics and grown for 6 h at the same conditions as the overnight culture. Appropriate concentrations of anhydrotetracycline (Sigma) were added to culture media as indicated. **Figure 4b**: 20 μL of each overnight culture were added to 980 μL of M9 minimal media containing selective antibiotics and grown for 24 h at 37C. Periodic samples of 10-200 μL of culture were collected for characterization by bulk fluorescence measurements. **For all bulk fluorescence measurements**: 10-200 μL of sampled culture were transferred to a 96-well plate (Costar) containing 0-190 μL of phosphate buffered saline (PBS). Fluorescence (FL) and optical density (OD) at 600 nm were then measured using a Synergy H1 plate reader (Biotek). The following settings were used: mCherry fluorescence (560 nm excitation, 630 nm emission).

### Bulk fluorescence data analysis

On each 96-well block there were two sets of controls; a media blank and *E. coli* Tax1 cells transformed with combination of control plasmids JBL002 and JBL644 (blank cells) and thus not expressing mCherry (**Supplementary Table 1**). The block contained three replicates of each control. OD and FL values for each colony were first corrected by subtracting the corresponding mean values of the media blank. The ratio of FL to OD (FL/OD) was then calculated for each well (grown from a single colony) and the mean FL/OD of blank cells was subtracted from each colony’s FL/OD value. Three biological replicates were collected from independent transformations, with three colonies characterized per transformation (9 colonies total). Occasional wells were discarded due to poor growth (OD < 0.1 at measurement), however, all samples contained at least 7 replicates over the three experiments. Means of FL/OD were calculated over replicates and error bars represent standard deviations (s.d).

### Small-scale “Hungate” fermentation

Small-scale fermentation assays were used to quantify oxygenated taxanes and taxadiene production in *E. coli* Tax1 or Tax1-QS. Experiments were performed with six biological replicates collected over three independent experiments (**Figure 2c, 2d**) or four biological replicates collected over two independent experiments (**Figure 3c, 3d, 3e, 3f, 4c, 4d**). For each experiment, plasmid combinations (SI Table 1) were transformed into chemically competent *E. coli* cells and plated on LB+Agar (Difco) plates containing appropriate antibiotics (100 μg/mL carbenicillin, 34 μg/mL chloramphenicol and/or 50 μg/mL spectinomycin). Plates were incubated approximately 17 hrs overnight at 30 °C. Individual colonies were inoculated into culture tubes containing LB and appropriate antibiotics and incubated at 30°C for roughly 16 hrs overnight to achieve an approximate OD600 of 3. For 2 mL batch fermentations, 50 μL of overnight cells were added to 1.95 mL of complete R-media (Supplementary Tables 5-7) and appropriate antibiotics in glass hungate tubes (ChemGlass). 0.1 mM IPTG was added for induction of the upstream pathway enzymes and p5Trc/p10Trc expression. 16-100 ng/mL aTc was added, as indicated, to induce P_*L,TetO1*_-STAR activated rSFPs. A 10% v/v dodecane layer (200 μL) was added in all fermentations. Hungate tubes were sealed with a rubber septa and plastic screw-cap (ChemGlass). PrecisionGlide 18G hypodermic needles (BD) were inserted into the rubber septa to allow for gas exchange. Hungate tubes were incubated at 22°C and 250 rpm for 96 hrs. After the fermentations were completed, the culture was centrifuged to collect the dodecane overlay. This overlay was subsequently diluted into hexane for analytical procedures described below.

### GC-MS analysis

Dodecane samples collected from batch fermentations were diluted at a ratio of 1:40 in n-hexane containing 5 mg/L ß-caryophyllene. The 5 mg/L ß-caryophyllene was utilized as a standard to calculate titer of taxadiene and oxygenated taxanes. GC-MS analysis was performed with an Agilent 7890 GC and Agilent HP-5ms-UI column (Ultra Inert, 30 m, 0.25 mm, 025 μm, 7 in cage). Helium was utilized as a carrier gas at a flow rate of 1 mL/min and the sample injection volume was 1 μL. The splitless method begins at 50 °C hold for 1 minute followed by a 10°C/min ramp to 200 °C and a final 5°C/min ramp to 270 °C. Mass spectroscopy data was collected for 22.5 minutes with an 11-minute solvent delay with an Agilent 7000 QQQ in scan mode using Electron Ionization (EI). m/z values ranging from 40-500 were scanned with a scan time of 528ms. MassHunter Workstation Qualitative Analysis software (vB.06.00) was utilized to integrate peaks on the chromatograms and determine their respective mass spectrums. The ratio of peak area of taxadiene (m/z 272) to the standard ß-caryophyllene (m/z 204) was used to calculate titer of taxadiene, while the ratio of the sum of all peaks of oxygenated taxanes (m/z 288) to ß-caryophyllene was used to calculate titer of the oxygenated taxanes. Means of titers were calculated over replicates and error bars represent s.d.

## Acknowledgements

The authors gratefully acknowledge Dr. Ryan Philippe for careful reading of the manuscript and the gift of *E. coli* Tax1 and plasmids p5Trc and p10Trc from Manus Bio. The pOSIP plasmid kit used for clonetegration was a gift from Drew Endy and Keith Shearwin (Addgene kit # 1000000035). This work was supported by an NSF CAREER award (1452441 to J. B. L.), an NSF CBET award (1803747 to J. B. L., K. T. and D. T.-E.), an NSF Graduate Research Fellowship (DGE-1144153 to C. J. G.), and an NIH Biotechnology Training Grant (T32-GM008449-23 to B. W. B.).

## Author Contributions

C.J.G., B.W.B., M.K.K., D.T.E, K.E.T., and J.B.L. designed the study. C.J.G., J.T.L., and J.A. performed experiments. All authors contributed to data analysis and preparation of the manuscript.

## Competing Financial Interests

A utility patent application has been filed for some of the developments contained in this article (US 5369-00519).

