## Supplementary Information for "Dynamic control of pathway expression with riboregulated switchable feedback promoters"

#### Table of Contents:


| | |
|---|---|
| Page 2-4 | Supplementary Table 1. Plasmids used in this study |
| Page 4-6 | Supplementary Table 2. Examples of DNA plasmid sequences |
| Page 6-8 | Supplementary Table 3. Sequence of Promoter and RBS variants |
| Page 9 | Supplementary Table 4. Strains used in this study |
| Page 9 | Supplementary Tables 5-7. Composition of R-media used for taxadiene fermentations |
| Page 10 | Supplementary Figure 1. Fold activation of rSFP variants |
| Page 10 | Supplementary Figure 2. Complete rSFP promoter screen |
| Page 11 | Supplementary Figure 3. Stress-response characterization of PmetN and PompF rSFPs |
| Page 12 | Supplementary Figure 4. Induction optimization of PmetN rSFPs |
| Page 12 | Supplementary Figure 5. Induction optimization of PompF rSFPs |
| Page 13 | Supplementary Figure 6. Example GC chromatogram for analysis of taxadiene and oxygenated taxane fermentations |
| Page 13 | Supplementary References |

### Supplementary Results

#### Supplementary Tables

**Supplementary Table 1. Plasmids used in this study.** apFAB parts were obtained from a previously published library of genetic parts[1]. Abbreviations are as follows: RBS 1 = ribosome binding site variant (see Supplementary Table 3), $P_R$ = tetR promoter[2], $P_{LTet,O1}$ = TetR repressible promoter[3], $P_{Lux}$ (BBa_R0062)* = LuxR inducible promoter[4], mCherry = red fluorescent protein, LuxR (BBa_C0062)* = AHL inducible transcription factor[4], tetR = tet repressor protein[3], TrrnB = rrnB terminator, BBa_B0015* = B0015 terminator, T500 = T500 terminator, dblTerm = dblTerm terminator, PJ2315 = BBa_J23115 promoter from the iGEM Registry of Standard Biological Parts (parts.igem.org), CmR = chloramphenicol resistance cassette, AmpR = ampicillin resistance cassette, SpcR = spectinomycin resistance cassette, p15A = p15A origin of replication, ColE1 = ColE1 origin of replication and CDF = CDF origin of replication.

| Plasmid # | Plasmid architecture | Name | Figure |
|---|---|---|---|
| pJBL002 | AmpR – ColE1 origin (Empty vector) | pJBL002 | 1d-e, 4b, SI Fig. 1, SI Fig. 3 |
| pJBL644 | SpcR – pCDF origin (Empty vecor) | pJBL644 | 1d-e, 4b, SI Fig. 1, SI Fig. 3 |
| pJBL6654 | $P_R$ – TetR – dblTerm – PL,TetO1 – STAR 8 – T500 – AmpR – ColE1 origin | $P_{L,TetO1}$-STAR | 1d-e, 2d, 3c-f, 4c-d, SI Fig. 1, SI Fig. 2, SI Fig. 3, SI Fig. 4, SI Fig 5 |
| pJBL6655 | PLux – STAR 8 – T500 – AmpR – ColE1 origin | $P_{Lux}$-STAR | 4b-c |
| pJBL6656 | PgntK – Target 8 – RBS 1 – mCherry – dblTerm – SpcR – pCDF origin | PgntK-mCherry | 1d, SI Fig. 1 |
| pJBL6657 | PompF – Target 8 – RBS 1 – mCherry – dblTerm – SpcR – pCDF origin | PompF-mCherry | 1d-e, 4b, , SI Fig. 1, SI Fig. 3 |
| pJBL6658 | PyeeF – Target 8 – RBS 1 – mCherry – dblTerm – SpcR – pCDF origin | PyeeF-mCherry | 1d-e, SI Fig. 1 |
| pJBL6659 | PompT – Target 8 – RBS 1 – mCherry – dblTerm – SpcR – pCDF origin | PompT-mCherry | 1d-e, SI Fig. 1 |
| pJBL6660 | PmetN – Target 8 – RBS 1 – mCherry – dblTerm – SpcR – pCDF origin | PmetN-mCherry | 1d-e, 4b, SI Fig. 1, SI Fig. 3 |
| pJBL6661 | Pb1762 – Target 8 – RBS 1 – mCherry – dblTerm – SpcR – pCDF origin | Pb1762-mCherry | 1d, SI Fig. 1 |
| pJBL6662 | PcarA – Target 8 – RBS 1 – mCherry – dblTerm – SpcR – pCDF origin | PcarA-mCherry | 1d, SI Fig. 1 |
| pJBL6663 | PfadL – Target 8 – RBS 1 – mCherry – dblTerm – SpcR – pCDF origin | PfadL-mCherry | 1d, SI Fig. 1 |
| pJBL6664 | PfecA – Target 8 – RBS 1 – mCherry – dblTerm – SpcR – pCDF origin | PfecA-mCherry | 1d, SI Fig. 1 |
| pJBL6665 | PuraA – Target 8 – RBS 1 – mCherry – dblTerm – SpcR – pCDF origin | PuraA-mCherry | 1d, SI Fig. 1 |

| | | | |
|---|---|---|---|
| pJBL6666 | PgrxA – Target 8 – RBS 1 – mCherry – dblTerm – SpcR – pCDF origin | PgrxA-mCherry | 1d, SI Fig. 1 |
| pJBL6667 | PmtgA – Target 8 – RBS 1 – mCherry – dblTerm – SpcR – pCDF origin | PmtgA-mCherry | 1d, SI Fig. 1 |
| pJBL6668 | PybcU – Target 8 – RBS 1 – mCherry – dblTerm – SpcR – pCDF origin | PycbU-mCherry | 1d, SI Fig. 1 |
| pJBL6669 | PycbS – Target 8 – RBS 1 – mCherry – dblTerm – SpcR – pCDF origin | PycbS-mCherry | 1d, SI Fig. 1 |
| pJBL6670 | PyhjX – Target 8 – RBS 1 – mCherry – dblTerm – SpcR – pCDF origin | PyhjX-mCherry | 1d, SI Fig. 1 |
| pJBL6671 | PatoA – Target 8 – RBS 1 – mCherry – dblTerm – SpcR – pCDF origin | PatoA-mCherry | 1d, SI Fig. 1 |
| pJBL6672 | Pb2970 – Target 8 – RBS 1 – mCherry – dblTerm – SpcR – pCDF origin | Pb2970-mCherry | 1d, SI Fig. 1 |
| pJBL6673 | PecpD – Target 8 – RBS 1 – mCherry – dblTerm – SpcR – pCDF origin | PecpD-mCherry | 1d, SI Fig. 1 |
| pJBL6674 | PgntK – Target 8 – RBS 1 – CYP725A4-tcCPR – dblTerm – CmR – p15a origin | PgntK-P450 | 2d, SI Fig 2 |
| pJBL6675 | PompF – Target 8 – RBS 1 – CYP725A4-tcCPR – dblTerm – CmR – P15a origin | PompF-P450 | 2d, 3d, 3f, 4d, SI Fig. 2, SI Fig. 5 |
| pJBL6676 | PyeeF – Target 8 – RBS 1 – CYP725A4-tcCPR – dblTerm – CmR – P15a origin | PyeeF-P450 | 2d, SI Fig 2 |
| pJBL6677 | PompT – Target 8 – RBS 1 – CYP725A4-tcCPR – dblTerm – CmR – P15a origin | PompT-P450 | 2d, SI Fig 2 |
| pJBL6678 | PmetN – Target 8 – RBS 1 – CYP725A4-tcCPR – dblTerm – CmR – P15a origin | PmetN-P450 | 2d, 3c, 3e, 4c, SI Fig. 2, SI Fig. 4 |
| pJBL6679 | Pb1762 – Target 8 – RBS 1 – CYP725A4-tcCPR – dblTerm – CmR – P15a origin | Pb1762-P450 | 2d, SI Fig 2 |
| pJBL6680 | PcarA – Target 8 – RBS 1 – CYP725A4-tcCPR – dblTerm – CmR – P15a origin | PcarA-P450 | 2d, SI Fig 2 |
| pJBL6681 | PfadL – Target 8 – RBS 1 – CYP725A4-tcCPR – dblTerm – CmR – P15a origin | PfadL-P450 | SI Fig 2 |
| pJBL6682 | PfecA – Target 8 – RBS 1 – CYP725A4-tcCPR – dblTerm – CmR – P15a origin | PfecA-P450 | SI Fig 2 |
| pJBL6683 | PuraA – Target 8 – RBS 1 – CYP725A4-tcCPR – dblTerm – CmR – P15a origin | PuraA-P450 | SI Fig 2 |
| pJBL6684 | PgrxA – Target 8 – RBS 1 – CYP725A4-tcCPR – dblTerm – CmR – P15a origin | PgrxA-P450 | SI Fig 2 |
| pJBL6685 | PmtgA – Target 8 – RBS 1 – CYP725A4-tcCPR – dblTerm – CmR – P15a origin | PmtgA-P450 | SI Fig 2 |
| pJBL6686 | PybcU – Target 8 – RBS 1 – CYP725A4-tcCPR – dblTerm – CmR – P15a origin | PybcU-P450 | SI Fig 2 |
| pJBL6687 | PycbS – Target 8 – RBS 1 – CYP725A4-tcCPR – dblTerm – CmR – P15a origin | PycbS-P450 | SI Fig 2 |
| pJBL6688 | PyhjX – Target 8 – RBS 1 – CYP725A4-tcCPR – dblTerm – CmR – P15a origin | PyhjX-P450 | SI Fig. 2 |
| pJBL6689 | PatoA – Target 8 – RBS 1 – CYP725A4-tcCPR – dblTerm – CmR – P15a origin | PatoA-P450 | SI Fig. 2 |
| pJBL6690 | PecpD – Target 8 – RBS 1 – CYP725A4-tcCPR – dblTerm – CmR – P15a origin | PecpD-P450 | SI Fig. 2 |

| | | | |
|---|---|---|---|
| N/A | PTrc – CYP725A4-tcCPR – rrnB – SpcR – SC101 | P5Trc | 2c |
| N/A | PTrc – CYP725A4-tcCPR – rrnB – CmR – p15a | P10Trc | 2c |
| pJBL6691 | apFAB346 – apFAB682 – EsaI -- LuxR — dblTerm – SpcR – SC101* | pQS | N/A |
| pJBL6692 | PJ23115 – Target 8 – RBS 1 – mCherry – dblTerm – SpcR – pCDF origin | PJ23115-mCherry | SI. Fig 3 |

**Supplementary Table 2. Examples of DNA plasmid sequences.** Abbreviations as described in Supplementary Table 1.

| Name | Sequence |
|---|---|
| PL,TetO1-STAR ($P_R$-TetR-dblTerm-PL,TetO1-STAR8-T500) | GAATTCTAAAGATCTTTTTCTCTATCACTGATAGGGAGTGGTAAAATAACTCTATCAACGAT<br>AGAGTGTCAACAAAAATTAGGAATTAATGATGTCGAGATTAGATAAAAGTAAAGTGATTAAC<br>AGCGCATTAGAGCTGCTTAATGAGGTCGGAATCGAAGGTTTAACAACCCGTAAACTCGCC<br>CAGAAGCTAGGTGTAGAGCAGCCTACATTGTATTGGCATGTAAAAAATAAGCGGGCTTTG<br>CTCGACGCCTTAGCCATTGAGATGTTAGATAGGCACCATACTCACTTTTGCCCTTTAGAAG<br>GGGAAAGCTGGCAAGATTTTTTACGTAATAACGCTAAAAGTTTTAGATGTGCTTTACTAAGT<br>CATCGCGATGGAGCAAAAGTACATTTAGGTACACGGCCTACAGAAAAACAGTATGAAACT<br>CTCGAAAATCAATTAGCCTTTTTATGCCAACAAGGTTTTTCACTAGAGAATGCATTATATGC<br>ACTCAGCGCTGTGGGGCATTTTACTTTAGGTTGCGTATTGGAAGATCAAGAGCATCAAGTC<br>GCTAAAGAAGAAAGGGAAACACCTACTACTGATAGTATGCCGCCATTATTACGACAAGCTA<br>TCGAATTATTTGATCACCAAGGTGCAGAGCCAGCCTTCTTATTCGGCCTTGAATTGATCAT<br>ATGCGGATTAGAAAAACAACTTAAATGTGAAAGTGGGTCTTAATAACACTGATAGTGCTAG<br>TGTAGATCACTACTAGAGCCAGGCATCAAATAAAACGAAAGGCTCAGTCGAAAGACTGGG<br>CCTTTCGTTTTATCTGTTGTTTGTCGGTGAACGCTCTCTACTAGAGTCACACTGGCTCACC<br>TTCGGGTGGGCCTTTCTGCGTTTATATACTAGAGTCCCTATCAGTGATAGAGATTGACATC<br>CCTATCAGTGATAGAGATACTGAGCACTGAACTGTATACATTCCCCGCAGGATAAGAGTAA<br>GTGAGAGTAGGTAGAGATTGAGGATGGGGATCTCAAAGCCCGCCGAAAGGCGGGCTTTT<br>TTTTGGATCCTTACTCGAGTCTAGACTGCAGGCTTCCTC |
| PLux-STAR (PLux-STAR8-T500) | CTAAAGATCTATATACTAGAGACCTGTAGGATCGTACAGGTTTACGCAAGAAAATGGTTTG<br>TTATAGTCGAATAAATGAACTGTATACATTCCCCGCAGGATAAGAGTAAGTGAGAGTAGGT<br>AGAGATTGAGGATGGGGATCTCAAAGCCCGCCGAAAGGCGGGCTTTTTTTTGGATCCTTA<br>CTCGAGTCTAGACTGCAGGCTTCCTC |
| Example rSFP Plasmid (PompF-TARGET 8-RBS 1-mCherry-dblTerm) | CGATCATCCTGTTACGGAATATTACATTGCAACATTTACGCGCAAAAACTAATCCGCATTCT<br>TATTGCGGATTAGTTTTTTCTTAGCTAATAGCACAATTTTCATACTATTTTTTGGCATTCTGG<br>ATGTCTGAAAGAAGATTTTGTGCCAGGTCGATAAAGTTTCCATCAGAAACAAAATTTCCGTT<br>TAGTTAATTTAAATATAAGGAAATCATATAAATAGATTAAAATTGCTGTAAATATCATCACGT<br>CTCTATGGAAATATGACGGTGTTCACAAAGTTCCTTAAATTTTACTTTTGGTTACATATTTTT<br>TCTTTTTGAAACCAAATCTTTATCTTTGTAGCACTTTCACGGTAGCGAAACGTTAGTTTGAA<br>TGGAAAGATGCCTGCAGACACATAAAGACACCAAACTCTCATCAATAGTTCCGTAAATTTT<br>TATTGACAGAACTTATTGACGGCAGTGGCAGGTGTCATAAAAAAAACCATGAGGGTAATAA<br>ATACCATCCTCAATCTCTACCTACTCTCACTTACTCTTATCCTGCGGGGAATGTATACAGTT<br>CATGTATATATTCCCCGCTTTTTTTTTGGATCTAGGAGGAAGGATCTATGGCGAGTAGCGA<br>AGACGTTATCAAAGAGTTCATGCGTTTCAAAGTTCGTATGGAAGGTTCCGTTAACGGTCAC<br>GAGTTCGAAATCGAAGGTGAAGGTGAAGGTCGTCCGTACGAAGGTACCCAGACCGCTAA<br>ACTGAAAGTTACCAAAGGTGGTCCGCTGCCGTTCGCTTGGGACATCCTGTCCCCGCAGTT<br>CCAGTACGGTTCCAAAGCTTACGTTAAACACCCGGCTGACATCCCGGACTACCTGAAACT<br>GTCCTTCCCGGAAGGTTTCAAATGGGAACGTGTTATGAACTTCGAAGACGGTGGTGTTGT<br>TACCGTTACCCAGGACTCCTCCCTGCAAGACGGTGAGTTCATCTACAAAGTTAAACTGCGT<br>GGTACCAACTTCCCGTCCGACGGTCCGGTTATGCAGAAAAAAACCATGGGTTGGGAAGCT<br>TCCACCGAACGTATGTACCCGGAAGACGGTGCTCTGAAAGGTGAAATCAAAATGCGTCTG<br>AAACTGAAAGACGGTGGTCACTACGACGCTGAAGTTAAAACCACCTACATGGCTAAAAAA<br>CCGGTTCAGCTGCCGGGTGCTTACAAAACCGACATCAAACTGGACATCACCTCCCACAAC<br>GAAGACTACACCATCGTTGAACAGTACGAACGTGCTGAAGGTCGTCACTCCACCGGTGCT<br>TAAGGATCCAAACTCGAGTAAGGATCTCCAGGCATCAAATAAAACGAAAGGCTCAGTCGA<br>AAGACTGGGCCTTTCGTTTTATCTGTTGTTTGTCGGTGAACGCTCTCTACTAGAGTCACAC<br>TGGCTCACCTTCGGGTGGGCCTTTCTGCGTTTATA |
| CYP725A4/tcCPR fusion | ATGGCTCTGTTATTAGCAGTTTTTTTTAGCATCGCTTTGAGTGCAATTGCCGGGATCTTGCT<br>GTTGCTCCTGCTGTTTCGCTCGAAACGTCATAGTAGCCTGAAATTACCTCCGGGCAAACT<br>GGGCATTCCGTTTATCGGTGAGTCCTTTATTTTTTTGCGCGCGCTGAGGAGCAATTCTCTG |

| | |
|---|---|
| | GAACAGTTCTTTGATGAACGTGTGAAGAAGTTCGGCCTGGTATTTAAAACGTCCCTTATCG<br>GTCACCCGACGGTTGTCCTGTGCGGGCCCGCAGGTAATCGCCTCATCCTGAGCAACGAA<br>GAAAAGCTGGTACAGATGTCCTGGCCGGCGCAGTTTATGAAGCTGATGGGAGAGAACTCA<br>GTTGCGACCCGCCGTGGTGAAGATCACATTGTTATGCGCTCCGCGTTGGCAGGCTTTTTC<br>GGCCCGGGAGCTCTGCAATCCTATATCGGCAAGATGAACACGGAAATCCAAAGCCATATT<br>AATGAAAAGTGGAAAGGGAAGGACGAGGTTAATGTCTTACCCCTGGTGCGGGAACTGGTT<br>TTTAACATCAGCGCTATTCTGTTCTTTAACATTTACGATAAGCAGGAACAAGACCGTCTGCA<br>CAAGTTGTTAGAAACCATTCTGGTAGGCTCGTTTGCCTTACCAATTGATTTACCGGGTTTC<br>GGGTTTCACCGCGCTTTACAAGGTCGTGCAAAACTCAATAAAATCATGTTGTCGCTTATTA<br>AAAAACGTAAAGAGGACTTACAGTCGGGATCGGCCACCGCGACGCAGGACCTGTTGTCT<br>GTGCTTCTGACTTTCCGTGATGATAAGGGCACCCCGTTAACCAATGACGAAATCCTGGAC<br>AACTTTAGCTCACTGCTTCACGCCTCTTACGACACCACGACTAGTCCAATGGCTCTGATTT<br>TCAAATTACTGTCAAGTAACCCTGAATGCTATCAGAAAGTCGTGCAAGAGCAACTCGAGAT<br>TCTGAGCAATAAGGAAGAAGGTGAAGAAATTACCTGGAAAGATCTTAAGGCCATGAAATAC<br>ACGTGGCAGGTTGCGCAGGAGACACTTCGCATGTTTCCACCGGTGTTCGGGACCTTCCG<br>CAAAGCGATCACGGATATTCAGTATGACGGATACACAATCCCGAAAGGTTGGAAACTGTT<br>GTGGACTACCTATAGCACTCATCCTAAGGACCTTTACTTCAACGAACCGGAGAAATTTATG<br>CCTAGTCGTTTCGATCAGGAAGGCAAACATGTTGCGCCCTATACCTTCCTGCCCTTTGGA<br>GGCGGTCAGCGGAGTTGTGTGGGTTGGGAGTTCTCTAAGATGGAGATTCTCCTCTTCGTG<br>CATCATTTCGTGAAAACATTTTCGAGCTATACCCCGGTCGATCCCGATGAAAAAATTTCCG<br>GCGATCCACTGCCGCCGTTACCGAGCAAAGGGTTTTCAATCAAACTGTTCCCTCGTCCGg<br>gcagcaccggatccCGCCGTGGTGGAAGTGATACACAGAAGCCCGCCGTACGTCCCACACCTC<br>TTGTTAAAGAAGAGGACGAAGAAGAAGAAGATGATAGCGCCAAGAAAAAGGTCACAATAT<br>TTTTTGGCACCCAGACCGGCACCGCCGAAGGTTTCGCAAAGGCCTTAGCTGAGGAAGCA<br>AAGGCACGTTATGAAAAGGCGGTATTTAAAGTCGTGGATTTGGATAACTATGCAGCGGAT<br>GACGAACAGTACGAAGAGAAGTTGAAAAAGGAAAAGCTAGCGTTCTTCATGCTCGCCACC<br>TACGGTGACGGCGAACCGACTGATAATGCCGCTCGCTTTTATAAATGGTTTCTCGAGGGT<br>AAAGAGCGCGAGCCATGGTTGTCAGATCTGACTTATGGCGTGTTTGGCTTAGGTAACCGT<br>CAGTATGAACACTTTAACAAGGTCGCGAAAGCGGTGGACGAAGTGCTCATTGAACAAGGC<br>GCCAAACGTCTGGTACCGGTAGGGCTTGGTGATGATGATCAGTGCATTGAGGACGACTTC<br>ACTGCCTGGAGAGAACAAGTGTGGCCTGAGCTGGATCAGCTCTTACGTGATGAAGATGAC<br>GAGCCGACGTCTGCGACCCCGTACACGGCGGCTATTCCAGAATACCGGGTGGAAATCTA<br>CGACTCAGTAGTGTCGGTCTATGAGGAAACCCATGCGCTGAAACAAAATGGACAAGCCGT<br>ATACGATATCCACCACCCGTGTCGCAGCAACGTGGCAGTACGTCGTGAGCTGCATACCCC<br>GCTGTCGGATCGTAGTTGTATTCATCTGGAATTCGATATTAGTGATACTGGGTTAATCTAT<br>GAGACGGGCGACCACGTTGGAGTTCATACCGAGAATTCAATTGAAACCGTGGAAGAAGCA<br>GCTAAACTGTTAGGTTACCAACTGGATACAATCTTCAGCGTGCATGGGGACAAGGAAGAT<br>GGAACACCATTGGGCGGGAGTAGCCTGCCACCGCCGTTTCCGGGGCCCTGCACGCTGC<br>GGACGGCGCTGGCACGTTACGCGGACCTGCTGAACCCTCCGCGCAAAGCCGCCTTCCTG<br>GCACTGGCCGCACACGCGTCAGATCCGGCTGAAGCTGAACGCCTTAAATTTCTCAGTTCT<br>CCAGCCGGAAAAGACGAATACTCACAGTGGGTCACTGCGTCCCAACGCAGCCTCCTCGA<br>GATTATGGCCGAATTCCCCAGCGCGAAACCGCCGCTGGGAGTGTTTTTCGCCGCAATAGC<br>GCCGCGCTTGCAACCTAGGTATTATAGCATCTCCTCCTCCCCGCGTTTCGCGCCGTCTCG<br>TATCCATGTAACGTGCGCGCTGGTCTATGGTCCTAGCCCTACGGGGCGTATTCATAAAGG<br>TGTGTGCAGCAACTGGATGAAGAATTCTTTGCCCTCCGAAGAAACCCACGATTGCAGCTG<br>GGCACCGGTCTTTGTGCGCCAGTCAAACTTTAAACTGCCCGCCGATTCGACGACGCCAAT<br>CGTGATGGTTGGACCTGGAACCGGCTTCGCTCCATTTCGCGGCTTCCTTCAGGAACGCG<br>CAAAACTGCAGGAAGCGGGCGAAAAATTGGGCCCGGCAGTGCTGTTTTTTGGGTGCCGC<br>AACCGCCAGATGGATTACATCTATGAAGATGAGCTTAAGGGTTACGTTGAAAAAGGTATTC<br>TGACGAATCTGATCGTTGCATTTTCACGAGAAGGCGCCACCAAAGAGTATGTTCAGCACA<br>AGATGTTAGAGAAAGCCTCCGACACGTGGTCTTTAATCGCCCAGGGTGGTTATCTGTATG<br>TTTGCGGTGATGCGAAGGGTATGGCCAGAGACGTACATCGCACCCTGCATACAATCGTTC<br>AGGAACAAGAATCCGTAGACTCGTCAAAAGCGGAGTTTTTAGTCAAAAAGCTGCAAATGG<br>ATGGACGCTACTTACGGGATATTTGGTAA |
| QS operon (apFAB346-apFAB682-EsaI-B0034-LuxR-dblTerm) | TTGACAATTAATCATCCGGCTCGTAATGTTTGTGGAGGGCCCAAGTTCACTTAAAAAGGAG<br>ATCAACAATGAAAGCAATTTTCGTACTGAAACATCTTAATCATGCTAAGGAGGTTTTCTAAT<br>GATGCTTGAACTGTTTGACGTCAGTTACGAAGAACTGCAAACCACCCGTTCAGAAGAACTT<br>TATAAACTTCGCAAGAAAACATTTAGCGATCGTCTGGGATGGGAAGTCATTTGCAGTCAGG<br>GAATGGAGTCCGATGAATTTGATGGGCCCGGTACACGTTATATTCTGGGAATCTGCGAAG<br>GACAATTAGTGTGCAGCGTACGTTTTACCAGCCTCGATCGTCCCAACATGATCACGCACA<br>CTTTTCAGCACTGCTTCAGTGATGTCACCCTGCCCGCCTATGGTACCGAATCCAGCCGTTT<br>TTTTGTCGACAAAGCCCGCGCACGTGCGCTGTTAGGTGAGCACTACCCTATCAGCCAGGT<br>CCTGTTTTTAGCGATGGTGAACTGGGCGCAAAATAATGCCTACGGCAATATCTATACGATT<br>GTCAGCCGCGCGATGTTGAAAATTCTCACTCGCTCTGGCTGGCAAATCAAAGTCATTAAA<br>GAGGCTTTCCTGACCGAAAAGGAACGTATCTATTTGCTGACGCTGCCAGCAGGTCAGGAT<br>GACAAGCAGCAACTCGGTGGTGATGTGGTGTCACGTACGGGCTGTCCGCCCGTCGCAGT<br>CACTACCTGGCCGCTGACGCTGCCGGTCTGATACTAGAGAAAGAGGAGAAATACTAGATG<br>AAAAACATAAATGCCGACGACACATACAGAATAATTAATAAAATTAAAGCTTGTAGAAGCAA<br>TAATGATATTAATCAATGCTTATCTGATATGACTAAAATGGTACATTGTGAATATTATTTACT<br>CGCGATCATTTATCCTCATTCTATGGTTAAATCTGATATTTCAATCCTAGATAATTACCCTAA |

| | |
|---|---|
| | AAAATGGAGGCAATATTATGATGACGCTAATTTAATAAAATATGATCCTATAGTAGATTATT<br>CTAACTCCAATCATTCACCAATTAATTGGAATATATTTGAAAACAATGCTGTAAATAAAAAAT<br>CTCCAAATGTAATTAAAGAAGCGAAAACATCAGGTCTTATCACTGGGTTTAGTTTCCCTATT<br>CATACGGCTAACAATGGCTTCGGAATGCTTAGTTTTGCACATTCAGAAAAAGACAACTATA<br>TAGATAGTTTATTTTTACATGCGTGTATGAACATACCATTAATTGTTCCTTCTCTAGTTGATA<br>ATTATCGAAAAATAAATATAGCAAATAATAAATCAAACAACGATTTAACCAAAAGAGAAAAA<br>GAATGTTTAGCGTGGGCATGCGAAGGAAAAAGCTCTTGGGATATTTCAAAAATATTAGGTT<br>GCAGTGAGCGTACTGTCACTTTCCATTTAACCAATGCGCAAATGAAACTCAATACAACAAA<br>CCGCTGCCAAAGTATTTCTAAAGCAATTTTAACAGGAGCAATTGATTGCCCATACTTTAAAA<br>ATTAATCTAGAGGATCCAAACTCGAGTAAGGATCTCCAGGCATCAAATAAAACGAAAGGCT<br>CAGTCGAAAGACTGGGCCTTTCGTTTTATCTGTTGTTTGTCGGTGAACGCTCTCTACTAGA<br>GTCACACTGGCTCACCTTCGGGTGGGCCTTTCTGCGTTTATACCTAGG |

**Supplementary Table 3. Sequence of Promoter and RBS variants**. Pstress promoters were PCR amplified from the *E. coli* K-12 MG1655 genome.

| Name | Sequence |
|---|---|
| RBS 1 | AGGAGGAA |
| B0034 RBS | AAGAGGAGAAA |
| $P_{L,TetO1}$ | TCCCTATCAGTGATAGAGATTGACATCCCTATCAGTGATAGAGATACTGAG<br>CAC |
| $P_{Lux}$ | ACCTGTAGGATCGTACAGGTTTACGCAAGAAAATGGTTTGTTATAGTCGAA<br>TAAA |
| PJ23115 | TTTATAGCTAGCTCAGCCCTTGGTACAATGCTAGC |
| PgntK | AATCTGTGACACCGAAAATGTTAGATTTAGGTTTCACCTTGTCACCGGGCG<br>GATCTATTTAAGCCCACAAATTTGAAGTAGCTCACACTTATACACTTAAGG<br>CATGGATGGATATTGCTTCTGATATTGTCCGGCTGGACAATGTTACCGATA<br>ACAGTTACCCGTAACATTTTTAATTCTTGTATTGTGGGGGCACCACT |
| PompF | CGATCATCCTGTTACGGAATATTACATTGCAACATTTACGCGCAAAAACTA<br>ATCCGCATTCTTATTGCGGATTAGTTTTTTCTTAGCTAATAGCACAATTTTC<br>ATACTATTTTTTGGCATTCTGGATGTCTGAAAGAAGATTTTGTGCCAGGTC<br>GATAAAGTTTCCATCAGAAACAAAATTTCCGTTTAGTTAATTTAAATATAAG<br>GAAATCATATAAATAGATTAAAATTGCTGTAAATATCATCACGTCTCTATGG<br>AAATATGACGGTGTTCACAAAGTTCCTTAAATTTTACTTTTGGTTACATATTT<br>TTTCTTTTTGAAACCAAATCTTTATCTTTGTAGCACTTTCACGGTAGCGAAA<br>CGTTAGTTTGAATGGAAAGATGCCTGCAGACACATAAAGACACCAAACTCT<br>CATCAATAGTTCCGTAAATTTTTATTGACAGAACTTATTGACGGCAGTGGC<br>AGGTGTCATAAAAAAAACCATGAGGGTAATAAATA |
| PyeeF | ATTAGCGGCCTCGGCTGCGGCTATTTACCCCGTTATCTGGCGCAACGTTT<br>TCTCGATAGTGGCGCGTTAATCGAGAAGAAAGTGGTCGCCCAAACTCTCT<br>TTGAACCCGTCTGGATTGGCTGGAACGAACAGACCGCAGGACTTGCCAGT<br>GGCTGGTGGCGGGATGAAATTTTAGCAAATAGTGCGATCGCCGGTGTTTA<br>TGCAAAATCTGATGACGGAAAATCAGCCATTTAAAGAAAAATTATTCTGAC<br>AAGCCTCTCATTCTCTTGTCATTTCCCCCCCATTTAGGCACAATGCGCCGC<br>TGTCAAAAAATGACTAAAAACCGACGTTTCATCAGCGTCGGTTATTTTTTG<br>CTTCAAACCAATCATTCATACCAAGAGGCCGGGCTTCGTACCGGATAGAT<br>ATTTACTAAAAATCGACAGTTGTTGTCGCTGAGGAATCCAAAAAAATGGGG<br>CAATTTTTTGCTTACGCGACGGTTATCACCGTAAAGGAGAATGACC |
| PompT | AACGGATAAGACGGGCATAAATGAGGAAGAAATGGCGCGCCCTGCAGGA<br>TTCGAACCTGCGGCCCACGACTTAGAAGTTCCTAGAACGACATTTTAAGTC<br>AACAACTTACCGCGCCATCTCTGCGCTCACACGTCCCACTACCTCAAAAC<br>ATGTAAAGCCTTGCAAGCCATTGCGAGGCCTTATGTGTCTCAGTTTTGTCC<br>CTCTTTTTTGTACTAAAAAACATAGTAATTGAGGATAAACCTCATGCTATTT<br>TCGCTTATATGCCTCTAAAGGCATGGCACTTAAATAGATAAAAGCACCACA<br>AAAGCATAAAAAAAACCACACAGTAAAACCGAAATATGAAACAATAACAGAT<br>AATTAAACCAAAAACAGATAGCGCATTGTGATAATCATTCAATACTAAACAA<br>AATATAAACAGTGGAGCAATATGTAATTGACTCATTAAGTTAGATATAAAAA<br>ATACATATTCAATCATTAAAACGATTGAATGGAGAACTTTT |
| PmetN | GGCGAAACTCTTCAACACTACCCTGCGGATGATGCGGGCAATAATAGATA<br>CCATCCAGATCGACATCTCGGTCCGCCAGCGACCAGTCCATCCACTCGGT<br>CAGCGTTTCAAACTGTGCTTCGGTAAATTTACCGCGAGCAATGCCAGACT<br>GGTTGGTTACTACCACCAGCGCAAAGCCCATTTTTTTTAGCTCGCGCATG<br>GCGTCAATAACACCGTCGATAAATTCAAAGTTGTCGATCTCATGGACATAG<br>CCGTGATCGACATTAATGGTGCCATCACGGTCAAGAAAAATTGCGGGTAC<br>GCTCTTCGCCACCTTTTATAGCTCCTTAATAAGGCATGTGACGCTAGTATC |

| | |
|---|---|
| | GCATGTTTCGACCTGCAAGAAAGTGCTCTTCGCATAAACCTGATTGATTTA<br>GACGTCTGGATGCCTTAACATCCATTTCATTGACGGCGTTGCCCGTTTCAG<br>GCATTCGAGATGCCACGACTAACTTAATGACGATAATAAATAATCA |
| Pb1762 | GATTATTGAACTGTTGTTCAAGCGTGGTTTCCTGACCAAAAAAGGGCGCTA<br>TATCCACTCCACCGACGCCGGAAAAGCGCTATTCCATTCGCTGCCGGAGA<br>TGGCGACGCGACCGGACATGACCGCGCACTGGGAATCGGTGCTGACGCA<br>AATCAGCGAAAAGCAGTGTCGCTATCAGGACTTTATGCAGCCGCTGGTGG<br>GGACGCTATATCAGCTTATTGATCAAGCCAAACGTACGCCGGTGCGGCAG<br>TTTCGCGGCATTGTGGCTCCGGGCAGTGGTGGCAGTGCTGATAAGAAAAA<br>GGCTGCACCGCGTAAACGTAGTGCGAAAAAAAGTCCGCCAGCAGATGAA<br>GTCGGAAGCGGGGCGATAGCGTAAGCGAGTGAATCTTTCGTGCTATTCGA<br>GTCATATTCTGAAATATCCAGCGGATCAAGAAAATTCGTTGGATATTTTTTT<br>TGCATGGATAAAATTATCGCCTCTAAAGTATGTAATAACAGGGAATGTG |
| PcarA | GGTCTTTTTGATATGCGAGATGTACTTGATCTCAATAATTTGTAACCACAAA<br>ATATTTGTTATGGTGCAAAAATAACACATTTAATTTATTGATTATAAAGGGC<br>TTTAATTTTTGGCCCTTTTATTTTTGGTGTTATGTTTTTAAATTGTCTATAAG<br>TGCCAAAAATTACATGTTTTGTCTTCTGTTTTTGTTGTTTTAATGTAAATTTT<br>GACCATTTGGTCCACTTTTTTCTGCTCGTTTTTATTTCATGCAATCTTCTTG<br>CTGCGCAAGCGTTTTCCAGAACAGGTTAGATGATCTTTTTGTCGCTTAATG<br>CCTGTAAAACATGCATGAGCCACAAAATAATATAAAAAATCCCGCCATTAA<br>GTTGACTTTTAGCGCCCATATCTCCAGAATGCCGCCGTTTGCCAGAAATTC<br>GTCGGTAAGCAGATTTGCATTGATTTACGTCATCATTGTGAATTAATATGCA<br>AATAAAGTGAGTGAATATTCTCTGGAGGGTGTT |
| PfadL | CGTTTGCTCTGCTTCTGCGCGGTTTGCAAACACGCGGCTGTAAGACGCGG<br>TGCAGTCGGAGTTGTCCATAATGGTGCCAACATCCATACAGCAGCAAACC<br>GGGGTTTCATCAGCACTACATTTACTCATCGTTGATTTCCTCTGTATGTGC<br>ACCCAAGGTGCCAGATAAACGTTGTGGATATTTTACGCTTCCGGAAAGTG<br>CTGCTCCAGTTGTTAATTCTGCAAAATCGGATAAGTGACCGAAATCACACT<br>TAAAAATGATCTAAAACAAAATTCACCCGAATCCATGAGTGCGCCACCTCC<br>AAATTTTGCCAGCTGGATCGCGTTTCTTAGATCATATTTGAAAAAAGATAGA<br>AACATACTTGCAACATTCCAGCTGGTCCGACCTATACTCTCGCCACTGGTC<br>TGATTTCTAAGATGTACCTCAGACCCTACACTTCGCGCTCCTGTTACAGCA<br>CGTAACATAGTTTGTATAAAAATAAATCATTGAGGTTATGGTC |
| PuraA | TACCGTTGTGCCAATTCTGCGTGCGGGTCTTGGTATGATGGACGGTGTGC<br>TGGAAAACGTTCCGAGCGCGCGCATCAGCGTTGTCGGTATGTACCGTAAT<br>GAAGAAACGCTGGAGCCGGTACCGTACTTCCAGAAACTGGTTTCTAACAT<br>CGATGAGCGTATGGCGCTGATCGTTGACCCAATGCTGGCAACCGGTGGTT<br>CCGTTATCGCGACCATCGACCTGCTGAAAAAAGCGGGCTGCAGCAGCATC<br>AAAGTTCTGGTGCTGGTAGCTGCGCCAGAAGGTATCGCTGCGCTGGAAAA<br>AGCGCACCCGGACGTCGAACTGTATACCGCATCGATTGATCAGGGACTGA<br>ACGAGCACGGATACATTATTCCGGGCCTCGGCGATGCCGGTGACAAAATC<br>TTTGGTACGAAATAAAGAATAAAAATAATTAAAGCCGACTTTAAGAGTCGG<br>CTTTTTTTTGAGTAAAGCGCCTATAACACATAATACAGAGGATAATACT |
| PgrxA | CGGAAATGGGTTCATCAGTGAAATGGCGAATGGAGCGATGGCCACAAATA<br>AGTTCAATGGTTGGCGTCATTATCTTTTTCTCTTTCTGAACGTGAATATTGC<br>GGTGGACGGTTCATCAGCTGTGGGGCAAGACGTTTTGCCACCTGAAGAAT<br>AACCACCACCGCAGCGGGAAGCATGAGCAAAACACCGAGAAAAATCATCA<br>GAATCTGCACTTCTGGCCGAGAAAATGGCTCAGGCAGCGACAGGGAGTC<br>GCTTACCGACAGCAGCGCCACCGCCAGTAGCATCATTCCGATAAATTCCA<br>GTATCAACACGCCTTTAGGCAATTTACCGATCGCGCGCATACGCTTCCCT<br>CTGCAAAGTGAGCCTTCAGTCTAAAACTTTTCACTGTATTGTGTTTAACAGT<br>TATAGCTTTTAGCAATTAATGCAACAGGTTAAACCTACTTTCAGCGAATACA<br>TTTTAGCGTGATCATTACAGGCATAAATCTATGAGGAGAGAAATA |
| PmtgA | CATTTAACTGGCGAAGCGATGACGGAAACGCGCAATGTGCTGATTGAAGC<br>GGCACGAATAACGCGCGGTGAAATCCGTCCTCTGGCCCAGGCCGATGCC<br>GCTGAACTGGATGCGTTGATTGTGCCGGGGGGGTTTGGCGCGGCGAAGA<br>ATTTAAGCAATTTTGCCAGTCTTGGTAGCGAATGCACCGTTGACCGTGAAT<br>TAAAGGCGCTGGCACAAGCGATGCATCAGGCCGGAAAACCGCTTGGTTTT<br>ATGTGTATTGCCCCGGCGATGCTGCCGAAAATTTTCGATTTCCCGCTGCG<br>TTTGACCATCGGTACTGATATCGATACCGCAGAAGTGCTGGAAGAGATGG<br>GCGCGGAGCATGTGCCGTGTCCTGTCGATGATATCGTGGTTGATGAAGAC<br>AATAAGATTGTCACCACCCCAGCATATATGCTGGCGCAGAACATTGCAGA<br>AGCGGCGAGCGGCATTGATAAGCTGGTTTCCCGCGTGCTGGTTCTGGCT<br>GA |
| PybcU | TCAAGAAATACGCATCTTATAGAAACGTCCTATGATAGGTTGAAATCAAGA<br>GAAATCACATTTCAGCAATACAGGGAAAATCTTGCTAAAGCAGGAGTTTTC<br>CGATGGATTACAAATATCCACGAACATAAAAGATATTACTATACCTTTGATA<br>ATTCATTACTATTTACTGAGAGCATTCAGAACACTACACAAATCTTTCCACG<br>CTAAATCATAACGTCCGGTTTCTTCCGTGTCAGCACCGGGGTGTTGGCAT |

| | |
|---|---|
| | AATACAATACATGTACGCGCTAAACCCTGTGTGCATCGTTTTTAATTATTCC<br>CGGACACTCCCGCAGAGAAGTTCCCCGTCAGGGCTGTGGACATAGTTAAT<br>CCGGGAATACAATGACGATTCATCGCACCTGGCATACATTAATAAATATTA<br>ACAATATGAAATTTCAACTCATTGTTTAGGGTTTGTTTAATTTTCTACACATA<br>CGATTCTGCGAACTTCAAAAAGCATCGGGAATAACACC |
| PycbS | CAGACTTTATTATTACACCACCGCTATTTGTGCTGAATCCGGCAAATGAGA<br>ATCTGTTACGCATTATGTACATTGGAGCGCCGTTGGCGAAAGACAGAGAA<br>ACCCTTTTCTTCACTAGCGTACGGGCAGTCCCTTCAACAACGAAGCGGAA<br>AGAGGGAAATACCCTGAAGATTGCCACACAAAGCGTCATCAAACTTTTCTG<br>GCGACCAAAAGGTTTAGCGTATCCCTTAGGCGAGGCTCCGGCGAAACTG<br>CGTTGCACTTCGTCAGCTGACATGGTTACGGTCAGTAACCCAACACCTTAT<br>TTCATTACCCTGACAGACCTGAAAATAGGTGGAAAAGTAGTTAAAAATCAA<br>ATGATTTCCCCCTTTGATAAATACCAATTTTCTCTGCCAAAGGGGGCCAAA<br>AATAGCAGCGTAACGTATCGAACCATCAATGACTACGGGGCGGAAACGCC<br>GCAACTCAACTGTAAATCGTAAGCCGTCTTCAGTTAAGAGAGCGAG |
| PyhjX | TAGGCTGGCGTGTTGACTCCCGGCTTGGCGATCTCCGACCCTGGGCGCA<br>AATCAGCTATAACCAGCAATTTGGCGAGAATATCTGGAAGGCGCAATCAG<br>GCCTGAGCCGGATGACGGCGACAAACCAGAACGGCAACTGGCTGGATGT<br>CACCGTAGGCGCTGATATGTTGCTCAATCAAAATATTGCCGCCTATGCCG<br>CGCTAACTCAGGCAGAAAATACCACTAATAATAGCGACTATCTGTATACGA<br>TGGGGGTTAGCGCCAGATTTTAACGTAACAGTCACAATTGAAACCATTAAA<br>TAACAATAGTTGTGGCGATAGTGGGTGCTAACTTACCAAATAATAAATTTG<br>GTGAATAATTGTCGCGTCATTCATTCCTGAACTAAGGCATTTCATTCCGTT<br>CTGATGGCATTTCATGCCGTTTTTCCCCAGGCATAAAGTGCACTTCGTTAT<br>GGTTGTCGGCAGAGATTTTTCCTTTTTATTACTGCAGGAATACTGCC |
| PatoA | GCGTTTGTTGATACCGGCATCGGTCCGCTCATCGTCAATGGTCGAGTCCG<br>CAAAGTGATTGCTTCACATATCGGCACCAACCCGGAAACAGGTCGGCGCA<br>TGATATCTGGTGAGATGGACGTCGTTCTGGTGCCGCAAGGTACGCTAATC<br>GAGCAAATTCGCTGTGGTGGAGCTGGACTTGGTGGTTTTCTCACCCCAAC<br>GGGTGTCGGCACCGTCGTAGAGGAAGGCAAACAGACACTGACACTCGAC<br>GGTAAAACCTGGCTGCTCGAACGCCCACTGCGCGCCGACCTGGCGCTAA<br>TTCGCGCTCATCGTTGCGACACACTTGGCAACCTGACCTATCAACTTAGC<br>GCCCGCAACTTTAACCCCCTGATAGCCCTTGCGGCTGATATCACGCTGGT<br>AGAGCCAGATGAACTGGTCGAAACCGGCGAGCTGCAACCTGACCATATTG<br>TCACCCCTGGTGCCGTTATCGACCACATCATCGTTTCACAGGAGAGCAAA<br>TA |
| Pb2970 | GGATTATTAAGTGGCTGTGCCAGCCATAATGAAAATGCCAGTTTACTGGC<br>GAAAAAACAGGCGCAAAATATCAGCCAAAACCTGCCGATTAAATCTGCGG<br>GATATACCTTAGTGCTGGCGCAAAGTAGCGGCACAACGGTAAAAATGACC<br>ATTATCAGCGAAGCGGGTACACAAACCACGCAGACGCCTGACGCCTTTTT<br>AACCAGCTATCAACGACAAATGTGCGCTGACCCGACGGTGAAATTAATGA<br>TCACTGAGGGAATTAATTACAGCATAACGATTAATGATACACGTACAGGTA<br>ACCAGTATCAGCGGAAACTGGATCGTACCACCTGTGGAATAGTCAAAGCA<br>TAACGTCGGGTAGATATAAATTGGCGCGGGTTGTTTTTCGTGACGCACGA<br>ATTTATCTCATTCAATGGCTGACAAAAATTCGTCACACTCTTAACCAGAGAC<br>AATCTCTTAATACAGACAAAGAGCATCTGCGAAAAATTGCACGCGGG |
| PecpD | CGGGCTGGAGGACGACGGTCAGATCAGCGCCAAAATCAACGGGCGGATT<br>TTCCCGCTTAACGGCAAGCGTAACTATCTCCCGCTCTCTCCCTATGGAAG<br>ATATGAGGTGGAGTTACAGAACAGCAAAAACTCACTCGACAGTTACGATAT<br>CGTCAGCGGCCGCAAAAGTCGTCTGACTCTCTATCCAGGCAATGTCGCTG<br>TCATTGAGCCAGAGGTGAAGCAGATGGTTACCGTCTCCGGTCGTATCCGT<br>GCGGAAGACGGCACACTGCTGGCTAACGCACGGATTAACAACCATATCG<br>GCCGAACCCGAACCGATGAAAACGGCGAGTTTGTCATGGACGTGGATAA<br>GAAATACCCCACTATCGATTTTCGCTACAGTGGCAATAAAACCTGCGAAGT<br>GGCTCTGGAACTCAACCAGGCGCGCGGTGCCGTCTGGGTCGGTGATGTG<br>GTCTGCAGCGGCCTCTCATCGTGGGCGGCGGTGACGCAGACAGGAGAAG<br>AGA |
| PfecA | GGGACAGAATTTACCGTCCGCCAGCAGGATAATTTCACGCAGCTTGACGT<br>GCAGCAGCACGCTGTGGAAGTGCTTCTCGCCAGTGCCCCCGCGCAAAAA<br>CGCATCGTGAACGCTGGTGAAAGCCTGCAGTTCAGCGCCTCTGAGTTTGG<br>CGCAGTGAAACCGCTGGATGACGAGAGTACAAGCTGGACGAAGGACATC<br>CTGAGCTTCAGCGATAAACCGCTGGGTGAGGTGATAGCCACGCTAACCC<br>GTTACCGCAACGGCGTGCTGCGCTGCGATCCCGCCGTTGCCGGGCTGCG<br>CCTGAGCGGGACGTTCCCGCTGAAAAATACCGATGCGATCCTGAACGTTA<br>TCGCGCAAACGCTTCCCGTTAAAATTCAGTCTATTACGCGGTACTGGATAA<br>ACATTTCACCACTGTAAGGAAAATAATTCTTATTTCGATTGTCCTTTTTACC<br>CTTCTCGTTCGACTCATAGCTGAACACAACAAAAATGATGATGGGGAAGGT |

**Supplementary Table 4. Strains used in this study**. Strains containing genomic insertions were created using the clonetegration platform to integrate the inserts using the HK022 plasmid into the *attB* site of the *E. coli* genome[5]. Successful integrations were identified by antibiotic selection and colony PCR according to the published protocol.[5]

| Strain | Strain Information | Genomic Insertion |
|---|---|---|
| *E. coli* Tax1 | E. coli containing genome integrated pathway enzymes for taxadiene biosynthesis (gift from Manus Bio) | N/A |
| *E. coli* Tax1-QS | Derived from E. coli Tax1 | *attB::EsaI-LuxR(*apFAB346-apFAB382-EsaI-LuxR-dblTerm – KmR ) |

**Supplementary Table 5.** Basal R-media recipe, per liter. Adapted from Biggs et al., 2016.[6]

| Component | Final media concentration (g/l) |
|---|---|
| $KH_2PO_4$ | 13.3 |
| $(NH_4)_2HPO_4$ | 4 |
| Citric Acid Monohydrate | 1.7 |
| Yeast Extract | 5 |
| HEPES | 23.83 |

**Supplementary Table 6.** 1000x Trace Element (TE) solution, per liter. Adapted from Biggs et al., 2016.

| Component | Final media concentration (mg/l) |
|---|---|
| EDTA | 8.4 |
| $H_3BO_3$ | 3.0 |
| $Zn(CH_3COO)_2$ | 8.0 |
| $CoCl_2 \cdot 6H_2O$ | 4.6 |
| $CuCl2 \cdot 2H_2O$ | 1.9 |
| $MnCl_2 \cdot 4H_2O$ | 24.0 |
| $Na_2MoO_4 \cdot 2H_2O$ | 2.9 |

**Supplementary Table 7.** Complete R-media compositions utilized for hungate tube fermentations. Adapted from Biggs et al., 2016.

| Component | Amount (mL) |
|---|---|
| Basal R-media | 35 |
| 32% v/v Glycerol | 1.3 |
| 1 M $MgSO_4$ | 0.171 |
| 0.1 M Ferric Citrate | 0.0858 |
| 1000x TE Solution | 0.035 |
| 1000x Antibiotic | 0.035 |
| 1 M Thiamine HCl | 0.00047 |

**Supplementary Figure 1.** Fold activation (ON/OFF) of rSFP variants containing unique envelope stress-response promoters. Fluorescence characterization was performed on *E. coli* transformed with plasmids encoding each rSFP controlling mCherry expression and $P_{L,TetO1}$-STAR in the absence and presence of 100 ng/mL aTc. Data represent mean values in units of arbitrary fluorescence/optical density (FL/OD) and error bars represent s.d. of at least n = 7 biological replicates. Mean fold activation for each rSFP variant is indicated above each bar.

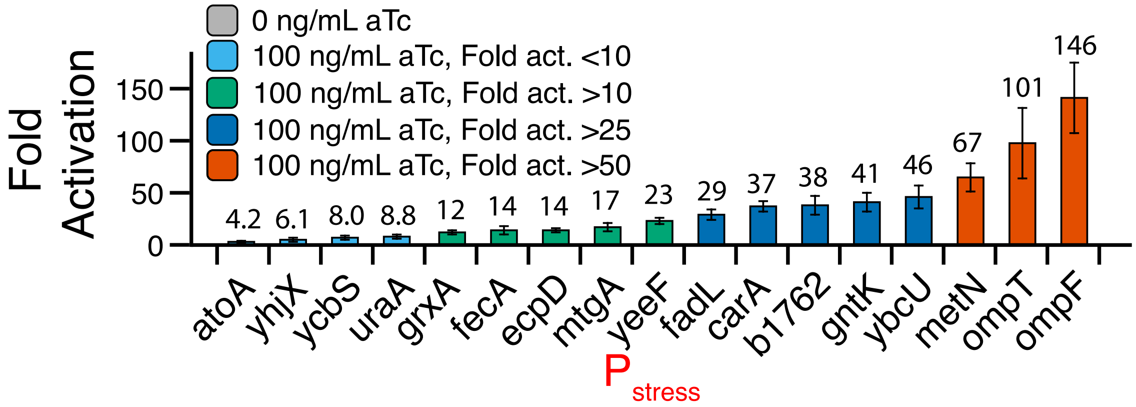


**Supplementary Figure 2.** Titers of fermentations after 96 hrs with *E. coli* Tax1 containing CYP725A4/tcCPR under control of complete rSFP library and $P_{L,TetO1}$-STAR with addition of 100 ng/mL aTc at inoculation. Dashed line represents production of oxygenated taxanes from p5Trc in Fig. 2C. Data represent mean values measured with GC-MS and error bars represent s.d. of at least n = 5 biological replicates.

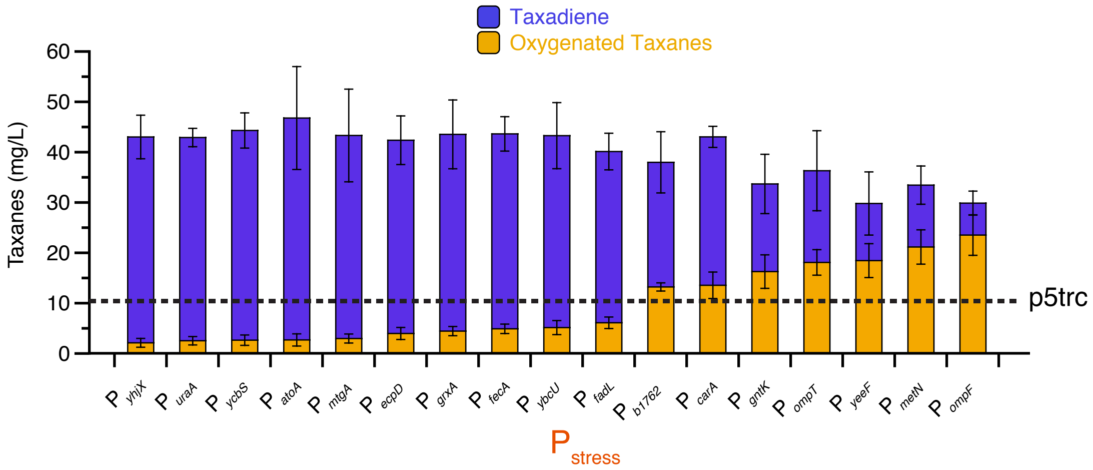

**Supplementary Figure 3.** Analysis of feedback-responsiveness of selected stress-response promoters to CYP725A4/tcCPR stress. (a) Schematic of plasmids used for fluorescence characterization of rSFP stress response. $P_{L,TetO1}$-STAR was used to activate expression from select rSFP plasmids. p10Trc was used to induce CYP275A4/tcCPR stress in comparison with an empty vector. Monitoring rSFP controlled expression of mCherry then allows the response to CYP275A4/tcCPR stress to be characterized. (b) Fluorescence characterization of cells containing select $P_{L,TetO1}$-STAR activated rSFPs controlling mCherry expression with 100 ng/mL aTc, and either an empty vector or the p10Trc vector to express CYP725A4/tcCPR and induce membrane stress. Fluorescence values were normalized to the empty vector control and error bars represent standard error of the mean. Experiments were performed as in main Figure 1d except 20 µL of each overnight culture were added to 490 µL of R-media containing selective antibiotics and grown for 4 h at 22C to closely mimic hungate fermentations. 100 ng/mL aTc was added after 4 hrs growth. After another 6 hrs of growth at 22C, 100 µL were sampled for characterization by bulk fluorescence measurements. Pcon = PJ23115. Data represent mean values of fermentation titers and error bars represent standard error of the mean (s.e.m) of at least n = 7 biological replicates.

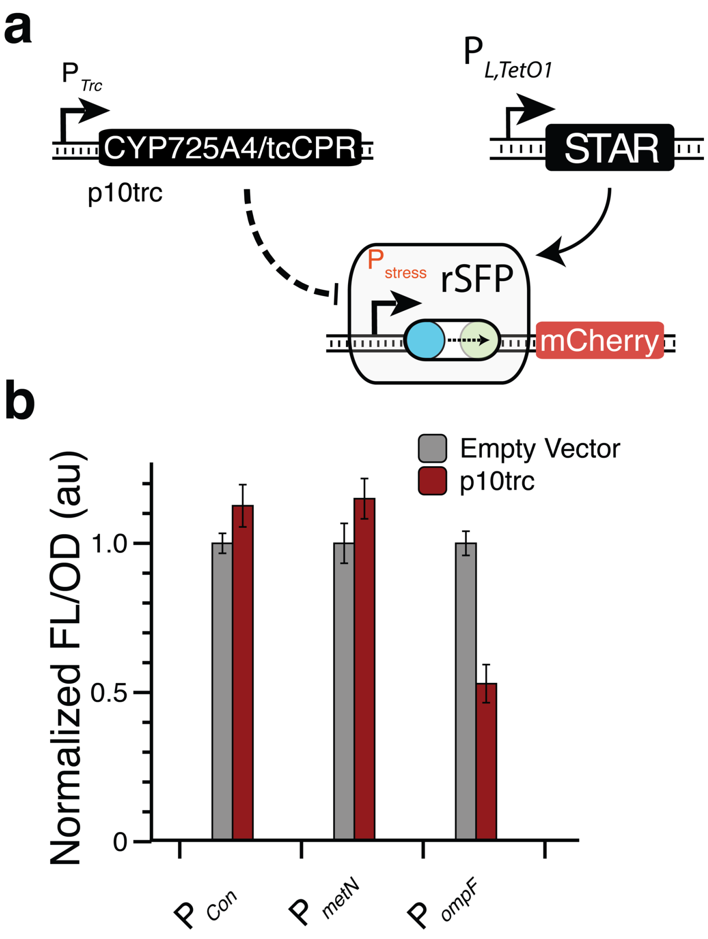

**Supplementary Figure 4.** Titers of fermentations after 96 hrs with *E. coli* Tax1 containing CYP725A4/tcCPR controlled by the $P_{metN}$ rSFP and $P_{L,TetO1}$-STAR under each induction condition. Dashed line represents production of oxygenated taxanes from p5Trc in Fig. 2C. Data represent mean values of fermentation titers and error bars represent s.d. of at least n = 3 biological replicates. Bold conditions indicate 100 ng/mL aTc induction at inoculation and the optimal inductions.

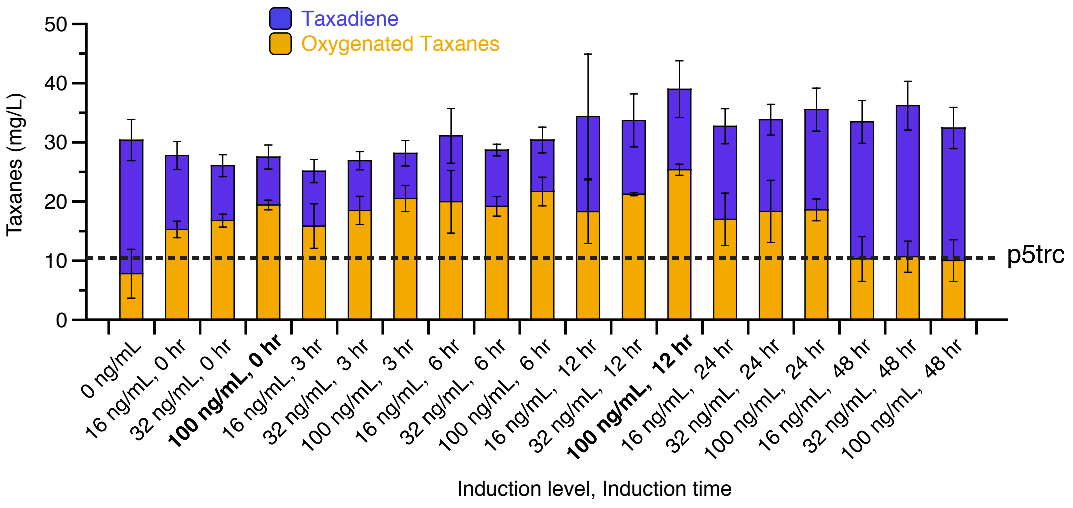


**Supplementary Figure 5.** Titers of fermentations after 96 hrs with *E. coli* Tax1 containing CYP725A4/tcCPR controlled by the $P_{ompF}$ rSFP and $P_{L,TetO1}$-STAR under each induction condition. Dashed line represents production of oxygenated taxanes from p5Trc in Fig. 2C. Data represent mean values of fermentation titers and error bars represent s.d. of at least n = 3 biological replicates. Bold conditions indicate 100 ng/mL aTc induction at inoculation and the optimal inductions.

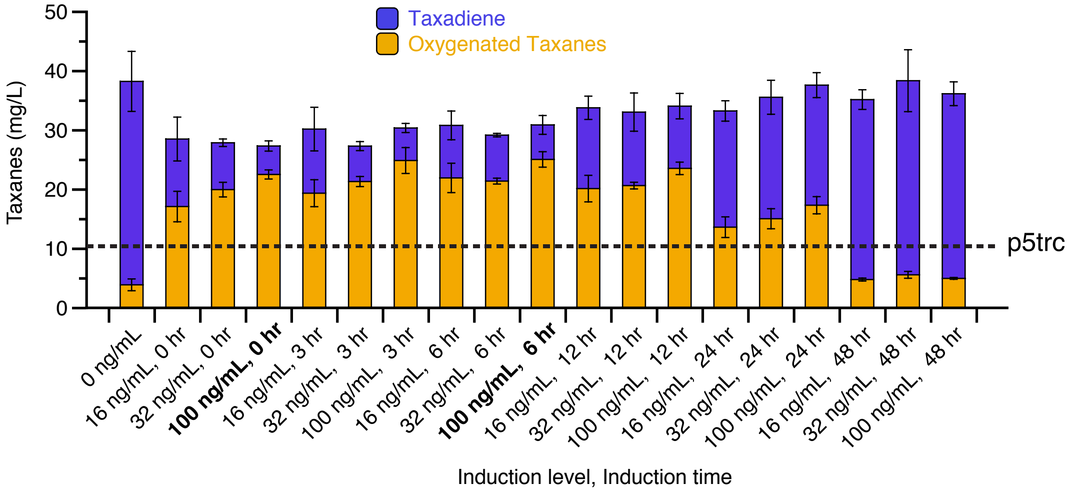

**Supplementary Figure 6.** Example GC chromatogram for analysis of taxadiene and oxygenated taxane fermentations. Taxadiene and oxygenated taxane peaks were previously described in Biggs et al.[6]

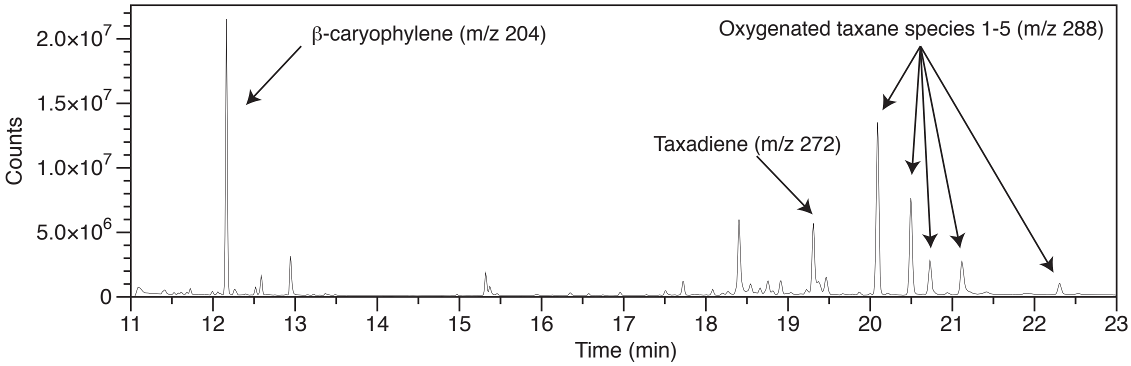
